# Evaluation of Cell Type Annotation R Packages on Single Cell RNA-seq Data

**DOI:** 10.1101/827139

**Authors:** Qianhui Huang, Yu Liu, Yuheng Du, Lana X. Garmire

## Abstract

Annotating cell types is a critical step in single cell RNA-Seq (scRNA-Seq) data analysis. Some supervised/semi-supervised classification methods have recently emerged to enable automated cell type identification. However, comprehensive evaluations of these methods are lacking to provide practical guidelines. Moreover, it is not clear whether some classification methods originally designed for analyzing other bulk omics data are adaptable to scRNA-Seq analysis. In this study, we evaluated ten cell-type annotation methods publicly available as R packages. Eight of them are popular methods developed specifically for single cell research (Seurat, scmap, SingleR, CHETAH, SingleCellNet, scID, Garnett, SCINA). The other two methods are repurposed from deconvoluting DNA methylation data: Linear Constrained Projection (CP) and Robust Partial Correlations (RPC). We conducted systematic comparisons on a wide variety of public scRNA-seq datasets as well as simulation data. We assessed the accuracy through intra-dataset and inter-dataset predictions, the robustness over practical challenges such as gene filtering, high similarity among cell types, and increased classification labels, as well as the capabilities on rare and unknown cell-type detection. Overall, methods such as Seurat, SingleR, CP, RPC and SingleCellNet performed well, with Seurat being the best at annotating major cell types. Also, Seurat, SingleR, CP and RPC are more robust against down-sampling. However, Seurat does have a major drawback at predicting rare cell populations, and it is suboptimal at differentiating cell types that are highly similar to each other, while SingleR and RPC are much better in these aspects. All the codes and data are available at: https://github.com/qianhuiSenn/scRNA_cell_deconv_benchmark.

## Introduction

Single cell RNA sequencing (scRNA-seq) has emerged as a powerful tool to enable the characterization of cell types and states in complex tissues and organisms at the single-cell level [1–5]. Annotating cell types amongst the cell clusters is a critical step before other downstream analyses, such as differential gene expression and pseudo time analysis [6–9]. Conventionally, a set of priorly known cell-type specific markers are used to label the cell types of the clusters manually. This process is laborious and often is a rate-limiting step for scRNA-seq analysis. This approach is also prone to bias and errors. The marker may not be specific enough to differentiate the cell subpopulations in the same dataset, or it may not be generic enough to be applied from one study to another. Automating the cell type labeling is critical to enhance reproducibility and consistency among single cell studies.

Recently some annotation methods have emerged to systematically assign cell types in the new scRNA-seq dataset, based on existing annotations from another dataset. Instead of using only top differentiating markers, most methods project or correlate the new cells onto similar cells in the well-annotated reference datasets, by leveraging the whole transcriptome profiles. These annotation methods are developed rapidly, at the same time benchmark datasets that the bioinformatics community agrees upon are lacking. These issues pose the urgent need to comprehensively evaluate these annotation methods using datasets with different biological variabilities, protocols and platforms. It is essential to provide practical guidelines for users. Additionally, identification of limitations of each method through comparisons will also help boost further algorithmic development, which in turn will benefit the scRNA-Seq community.

In this study, we evaluated ten cell annotation methods publicly available as R packages (**Table 1**). Eight of them are popular methods developed specifically for single cell research (Seurat [10], scmap [11], SingleR [12], CHETAH [13], SingleCellNet [14], scID [15], Garnett [16], SCINA [17]). Those methods can be further divided into two categories: Seurat, scmap, CHETAH, SingleCellNet, and scID utilize the gene expression profile as a reference without prior knowledge in signature sets, while Garnett and SCINA require additional pre-defined gene markers as the input. Additionally, to potentially leverage existing deconvolution methods for other bulk omics data, we also included two modified methods: Linear Constrained Projection (CP) and Robust Partial Correlations (RPC) that are popular in DNA methylation analysis [18]. We conducted systematic comparisons on six publicly available scRNA-seq datasets (**Table 2**) varying by species, tissue and sequencing protocol, as well as six sets of simulation data with known truth measure.

**Table 1.** List of Single-Cell RNA-sequencing/Methylation Cell Deconvolution tools benchmarked in this study.

| **Software** | **Method/Algorithm** | **Bulk/Single**<br>**Reference** | **Require Pre-**<br>**defined Marker**<br>**Genes** | **Allow Unknown** | **Version**<br>**Under R 3.6.0** | **Reference** |
| --- | --- | --- | --- | --- | --- | --- |
| SingleR | Correlation-based with Iterative Tuning | Bulk | No | No | SingleR_1.0.0 | [12] |
| CP | Reference-based method using Constrained Projection | Bulk | No | No | EpiDISH_2.0.2 | [18] |
| RPC | Reference-based Robust Partial Correlations | Bulk | No | No | EpiDISH_2.0.2 | [18] |
| Garnett | Elastic net Multinomial Regression | Single | Yes | Yes | garnett_0.1.4 | [16] |
| SCINA | Bimodal Distribution assumption for marker genes | Single | Yes | Yes | SCINA_1.1.0 | [17] |
| Seurat | Define anchor with CCA, L2-norm and MNN | Single | No | No | Seurat_3.0.1 | [10] |
| singleCellNet | Multi-Class Random Forest | Single | No | No | singleCellNet_0.1.0 | [14] |
| CHETAH | Correlation-based with Hierarchical Classification | Single | No | Yes | CHETAH_1.1.2 | [13] |
| scmap | K-nearest-neighbor classification<br>with cosine similarity | Single | No | Yes | scmap_1.6.0 | [11] |
| scID | Fisher's Linear Discriminant<br>Analysis-like Framework | Single | No | Yes | scID_0.0.0.9000 | [15] |

**Table 2.**
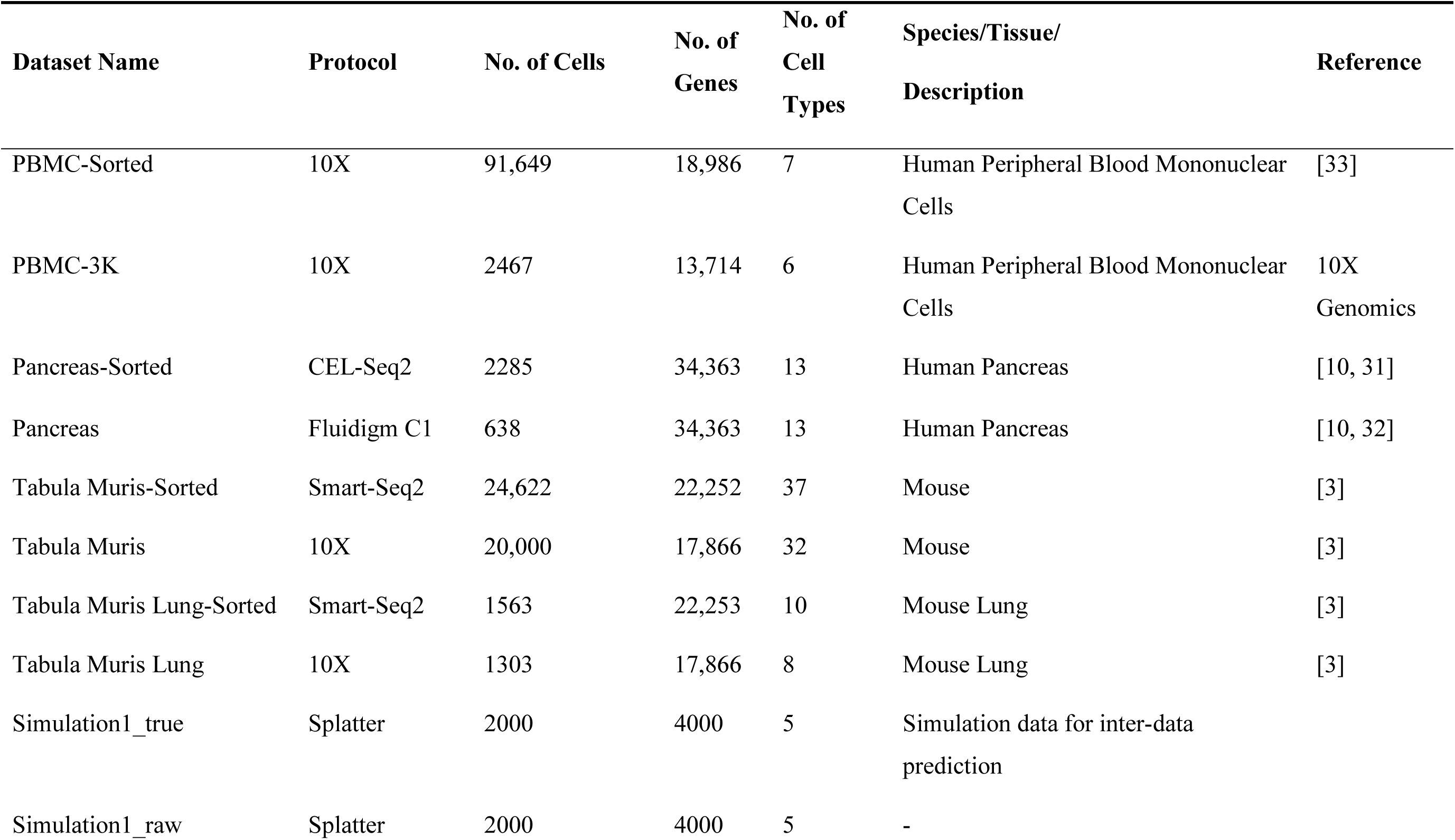
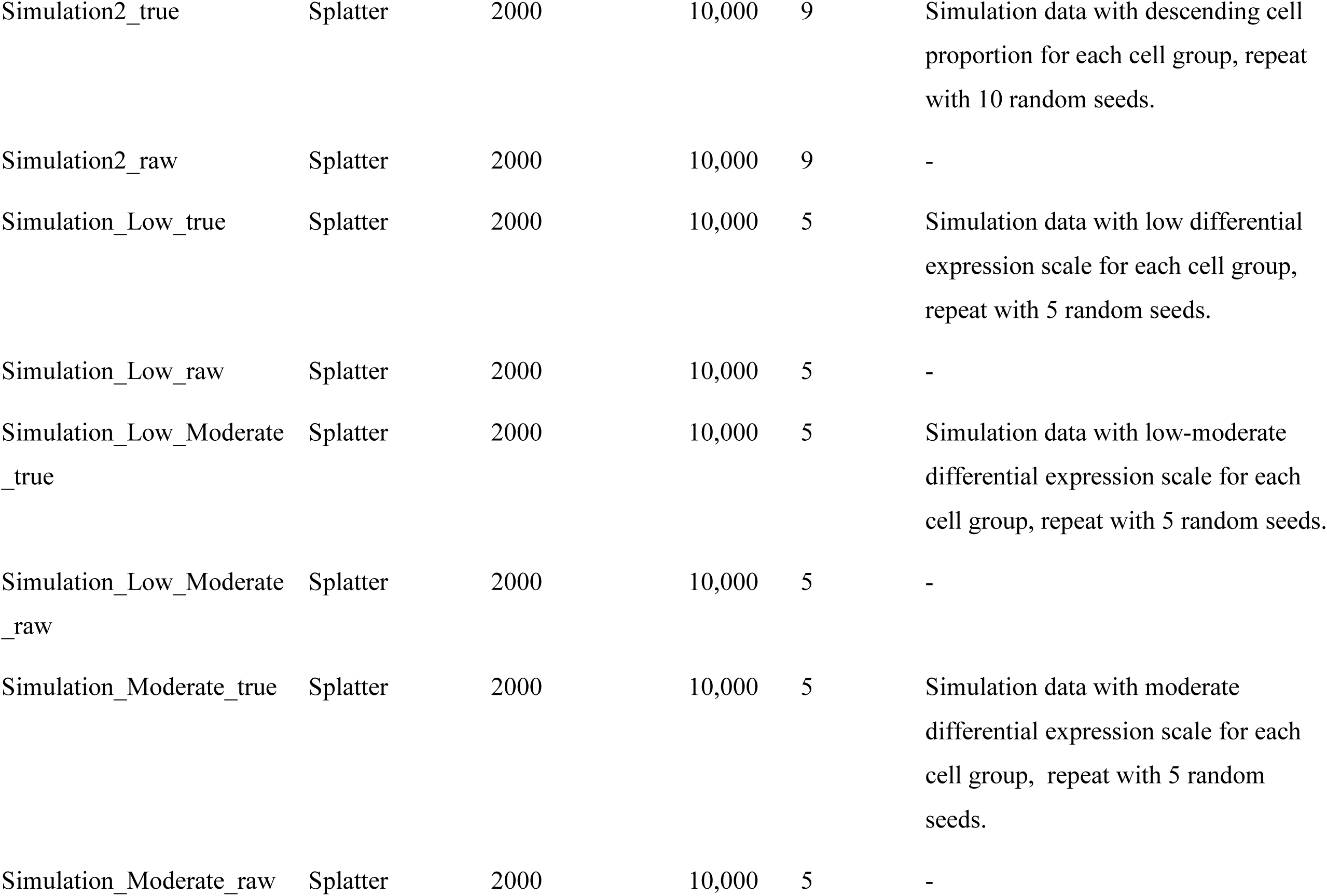
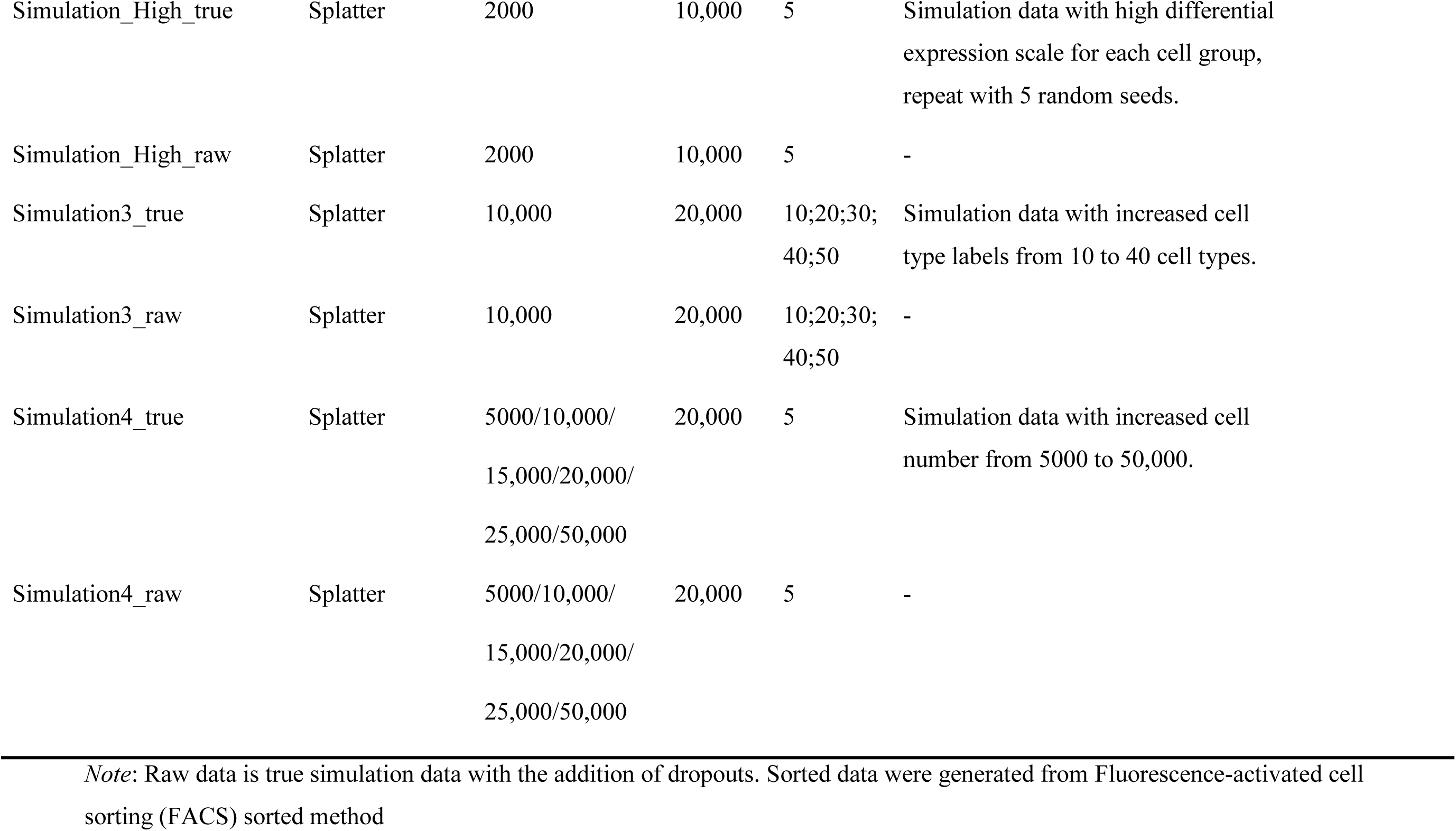
Datasets used in this study.

| **Dataset Name** | **Protocol** | **No. of Cells** | **No. of Genes** | **No. of Cell Types** | **Species/Tissue/Description** | **Reference** |
| --- | --- | --- | --- | --- | --- | --- |
| PBMC-Sorted | 10X | 91,649 | 18,986 | 7 | Human Peripheral Blood Mononuclear Cells | [33] |
| PBMC-3K | 10X | 2467 | 13,714 | 6 | Human Peripheral Blood Mononuclear Cells | 10X Genomics |
| Pancreas-Sorted | CEL-Seq2 | 2285 | 34,363 | 13 | Human Pancreas | [10, 31] |
| Pancreas | Fluidigm C1 | 638 | 34,363 | 13 | Human Pancreas | [10, 32] |
| Tabula Muris-Sorted | Smart-Seq2 | 24,622 | 22,252 | 37 | Mouse | [3] |
| Tabula Muris | 10X | 20,000 | 17,866 | 32 | Mouse | [3] |
| Tabula Muris Lung-Sorted | Smart-Seq2 | 1563 | 22,253 | 10 | Mouse Lung | [3] |
| Tabula Muris Lung | 10X | 1303 | 17,866 | 8 | Mouse Lung | [3] |
| Simulation1_true | Splatter | 2000 | 4000 | 5 | Simulation data for inter-data prediction |  |
| Simulation1_raw | Splatter | 2000 | 4000 | 5 | - |  |

|  |  |  |  |  |  |
| --- | --- | --- | --- | --- | --- |
| Simulation2_true | Splatter | 2000 | 10,000 | 9 | Simulation data with descending cell proportion for each cell group, repeat with 10 random seeds. |
| Simulation2_raw | Splatter | 2000 | 10,000 | 9 | - |
| Simulation_Low_true | Splatter | 2000 | 10,000 | 5 | Simulation data with low differential expression scale for each cell group, repeat with 5 random seeds. |
| Simulation_Low_raw | Splatter | 2000 | 10,000 | 5 | - |
| Simulation_Low_Moderate_true | Splatter | 2000 | 10,000 | 5 | Simulation data with low-moderate differential expression scale for each cell group, repeat with 5 random seeds. |
| Simulation_Low_Moderate_raw | Splatter | 2000 | 10,000 | 5 | - |
| Simulation_Moderate_true | Splatter | 2000 | 10,000 | 5 | Simulation data with moderate differential expression scale for each cell group, repeat with 5 random seeds. |
| Simulation_Moderate_raw | Splatter | 2000 | 10,000 | 5 | - |
| Simulation_High_true | Splatter | 2000 | 10,000 | 5 | Simulation data with high differential expression scale for each cell group, repeat with 5 random seeds. |
| Simulation_High_raw | Splatter | 2000 | 10,000 | 5 | - |
| Simulation3_true | Splatter | 10,000 | 20,000 | 10;20;30;40;50 | Simulation data with increased cell type labels from 10 to 40 cell types. |
| Simulation3_raw | Splatter | 10,000 | 20,000 | 10;20;30;40;50 | - |
| Simulation4_true | Splatter | 5000/10,000/<br>15,000/20,000/<br>25,000/50,000 | 20,000 | 5 | Simulation data with increased cell number from 5000 to 50,000. |
| Simulation4_raw | Splatter | 5000/10,000/<br>15,000/20,000/<br>25,000/50,000 | 20,000 | 5 | - |

*Note:* Raw data is true simulation data with the addition of dropouts. Sorted data were generated from Fluorescence-activated cell sorting (FACS) sorted method

## Results

### Intra-dataset accuracy evaluation

We first tested the classification accuracy of 10 methods (**Table 1**) on six publicly available scRNA-seq datasets (**Table 2**). These datasets include two human pancreatic islet datasets (GSE85241 and GSE86469), two whole mouse datasets (Tabula Muris, or TM-Full), and two peripheral blood mononuclear cells datasets (PBMC). Since Tabula Muris datasets are heterogeneous in terms of tissue contents, to evaluate the tools’ performance on homogeneous data, we down sampled them separately into two mouse lung datasets (Tabula Muris-Lung, or TM- Lung) by taking cells from lung tissue only. This results in eight real scRNA-seq datasets (**Table 1**). To avoid potential bias, we used the 5-fold cross validation scheme to measure the averaged accuracy in the 1-fold holdout subset. We used three different performance measurement metrics: overall accuracy, adjusted rand index (ARI), and V-measure [19–20] (see **Materials and methods**). The evaluation workflow is depicted in Figure S1.

**Figure 1A-C** show the classification accuracy metrics on eight datasets. The top-five performing annotation methods are Seurat, SingleR, CP, singleCellNet and RPC. Seurat has the best overall classification performance in the 5-fold cross validation evaluation. On average, the three accuracy metrics from Seurat are significantly higher (Wilcoxon paired rank test, P<0.05) than 9 other methods. SingleR has the second-best performance after Seurat, with all three metrics higher than 8 other methods, among which 6 pair-wise method comparisons achieved statistical significance (Wilcoxon paired rank test P < 0.05). Though slightly lower in average metric scores, the classification performance of both singleCellNet and CP are comparable to SingleR.

**Figure 1.**
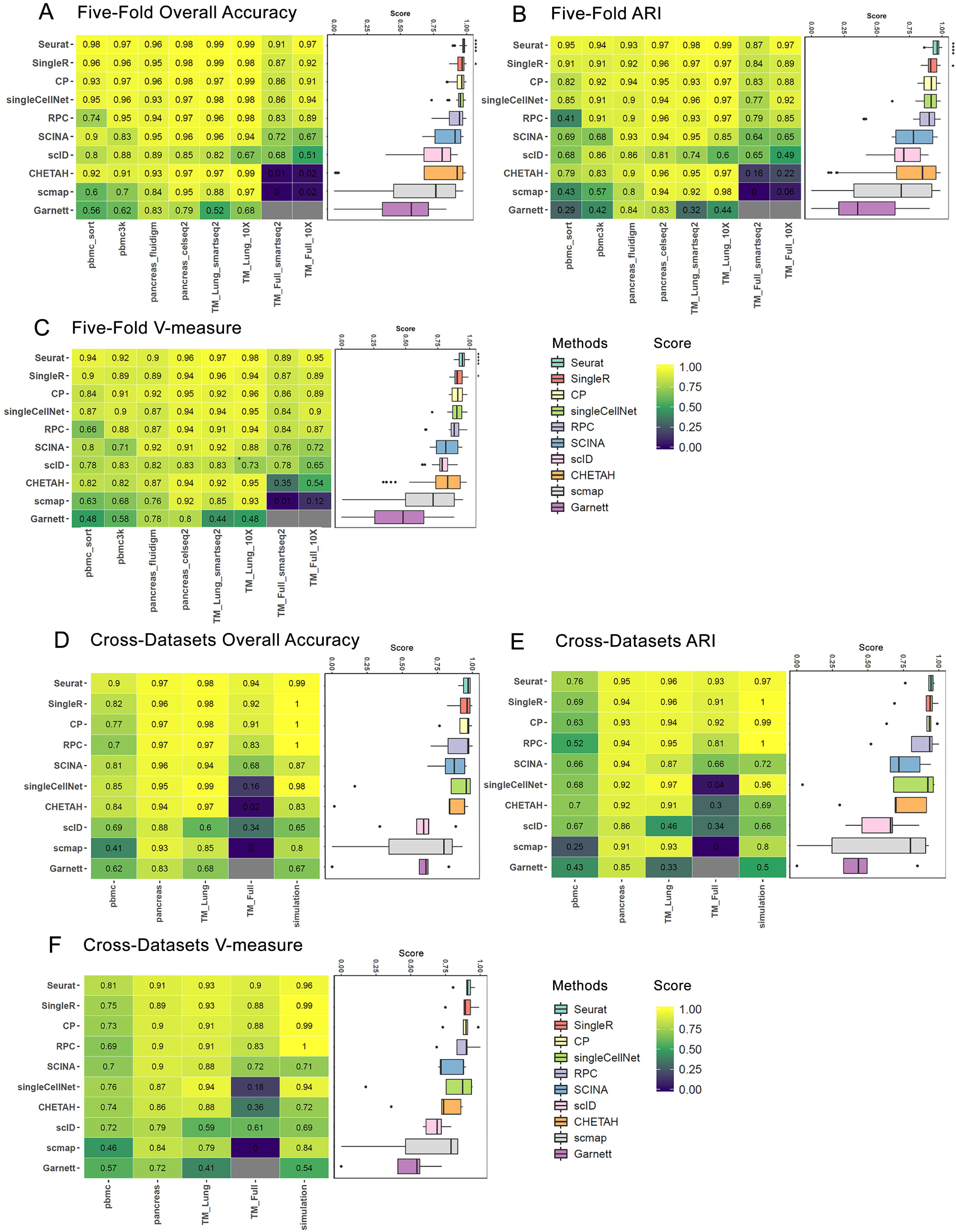
Inter-data and cross-date accuracy comparison. **(A-C)** Within data accuracy comparison, shown as heatmaps of three classification metrics, (**A**) overall accuracy, (**B**) adjusted rand index (ARI), and (**C**) v-measure across eight real datasets. For each dataset, a 5-fold cross validation is performed: using four folds as the reference and one-fold as the query. **(D-F)** Between-data accuracy comparison, shown as heatmaps of three classification metrics, (**D**) overall accuracy, (**E**) adjusted rand index (ARI), and (**F**) v-measure across four pairs of experimental datasets and one pair of simulation datasets. PBMC cell pair: PBMC-sorted-ref and PBMC-3K-query; pancreas pair: pancreas-celseq2-ref and pancreas-fluidigm-query; TM-Full pair: TM-Full-smartseq2-ref and TM-Full-10X-query; TM-Lung pair: TM-Lung-smartseq2-ref and TM-Lung-10x-query; simulation: true-ref and dropout-masked raw. TM-Lung datasets pair was down sampled from TM-Full datasets pair by taking cells from lung tissue only. Within the simulation datasets pair, the true assay without dropouts (true-ref) was used as the reference and the raw assay with dropout mask (raw-query) was used as the query. The columns are datasets, and the rows are annotation methods. The heatmap scale is shown on the figure, where the brighter yellow color indicates a better classification accuracy score. On the right of each heatmap is a boxplot to summarize the classification metrics among methods. Box colors represent different methods as shown in the figure. The methods in the heatmap and the boxplot are arranged in descending order by their average metrics score across all datasets. Some methods failed to produce a prediction for certain data sets (indicated by grey squares). ****: significantly higher (P<0.05) than 9 other methods using pairwise Wilcoxon test. ***: significantly higher (P<0.05) than 8 other methods using pairwise Wilcoxon test. **: significantly higher (P<0.05) than 7 other methods using pairwise Wilcoxon test. *: significantly higher (P<0.05) than 6 other methods using pairwise Wilcoxon test.

In order to test the influence of cell-type number on tool’s performance, we next evaluated the TM-Full and TM-Lung results. As shown in **Table 2**, for two TM-Full datasets from 10X and Smart-Seq2 platforms, which contain a large number of cell types (32 and 37 cell types, respectively), we took a subset of cells from the lung tissue and created two TM-Lung datasets that are relatively small in cell-type numbers, with 8 and 10 cell types for 10X and Smart-Seq2 platforms, respectively. Most methods perform well for both TM-Lung datasets with ARI > 0.9. However, some of the methods had a drop of performance on the two TM-Full datasets. The increased classification labels imposed a challenge. Garnett failed to predict on such large TM-Full datasets. Additionally, SCINA, CHETAH and scmap have significantly lower classification metrics on TM-Full datasets, compared to those on TM-Lung datasets. On the contrary, the previously mentioned top-five methods are more robust despite the increase of complexity in TM-Full datasets. Again, Seurat yields the best metric scores in both TM-Full datasets, demonstrating its capability at analyzing complex datasets.

### Cross-dataset accuracy evaluation

To evaluate the annotation tools in a more realistic setting, we conducted cross-dataset performance evaluation on 10 datasets (5 pairs), where the referencing labels were obtained from one dataset and the classification was done on another dataset of the same tissue type (**Table 2**). Within a pair, we used FACS-sorted, purified dataset as the reference data, and the remaining one as the query data (see **Materials and methods**). Among the 5 pairs of datasets, 4 are real experimental data: PBMC cell pair with PBMC-sorted-ref and PBMC-3K-query; pancreas cell pair with pancreas-celseq2-ref and pancreas-fluidigm-query; TM-Full pair with TM-Full-smartseq2- ref and TM-Full-10X-query; TM-Lung pair with TM-Lung-smartseq2-ref and TM-Lung-10x- query. The last pair is simulation datasets with the pre-defined truth, where the true assay without dropouts (simulation_true-ref) was used as the reference and the raw assay with dropout mask (simulation_raw-query) was used as the query.

**Figure 1D-F** shows the classification accuracy metrics on the above mentioned 5 pairs of query and reference datasets. The top 3 performing annotation methods in the descending rank order are Seurat, SingleR, and CP, the same as those in the same-dataset cross validation results (**Figure 1A-C**). In particular, they all perform very well on the simulation data with known truth measures, as all three accuracy metrics are above 0.96. RPC is ranked 4th, slightly better than SCINA. Similar to the 5-fold cross validation evaluation, methods such as scID, CHETAH, scmap and Garnett are persistently ranked among the lowest-performing methods for accuracy. Interestingly, singleCellNet, the method that performs relatively well (ranked 4th) in the same-dataset 5-fold cross validation, is now consistently ranked on the 6th, behind RPC and SCINA, due to drop of performance in TM-Full datasets. Besides annotation methods, the accuracy scores are also much dependent on the datasets. For example, on complex PBMC datasets, even Seurat only reaches 0.76 for ARI. A further examination of the confusion matrices (Figure S2) for Seurat, SingleR, CP, singleCellNet and RPC reveals that the challenge comes from distinguishing highly similar cell types such as CD4+ T cells vs. CD8+ T cells or Dendritic cells vs. CD14+ Monocytes in the PBMC datasets.

We also performed batch corrected cross-dataset accuracy evaluations on 4 pairs of experimental datasets. For each pair of data, both reference and query datasets were aligned using CCA [10,21]. The result is illustrated in Figure S3A-B. Most methods do not benefit from aligning and integrating the datasets (Figure S3B). None of the other methods exceeds the performance of Seurat in all three metrics after the batch correction (Figure S3A). The drop of performance in those methods may be attributed to the fact that aligned datasets contain negative values after the matrix correction and subtraction from the integration algorithm used in Seurat. In addition, some algorithms require non-normalized data matrix as the input, while batch-corrected matrix from Seurat is normalized, which may violate some models’ assumptions.

Altogether, these results from both experimental and simulation data indicate that Seurat has the best overall accuracy among the annotation methods in comparison, based on intra-dataset prediction and cross-dataset prediction [10].

### The effect of cell type similarity

Since it is challenging to distinguish highly similar cell populations using cross-data evaluation, we next conducted simulations. We designed 20 simulation data sets composed of five cell groups with varying levels of differential expression. Similar to others [22], we used Splatter [23] to predefine the same set of differential expression (DE) genes in simulation datasets, and only differed the magnitudes of DE, from low, low-moderate, moderate to high (**Figure 2A**). When cell populations are more separable, the classification task is easy for the majority of methods. As the cell populations become less separable, all methods show a decrease in their performance (**Figure 2B-D**). The degrees of such decrease vary among the methods though. SingleR, RPC, Seurat, singleCellNet and CP are in the first class that are relatively more robust than the other five methods. SingleR and RPC are ranked the 1st and 2nd for their robustness against cell type similarity, with all three metric scores above 0.9. Seurat is ranked the 4th after singleCelNet (the 3rd) when the samples are least separable (low DE), exposing its slight disadvantage. Garnett failed to predict when cell-cell similarity is high (low DE). In this context, the pre-defined marker genes may be ‘ambiguous’ to discriminate in multiple cell types, which may cause problems for Garnett to train the classifier.

**Figure 2.**
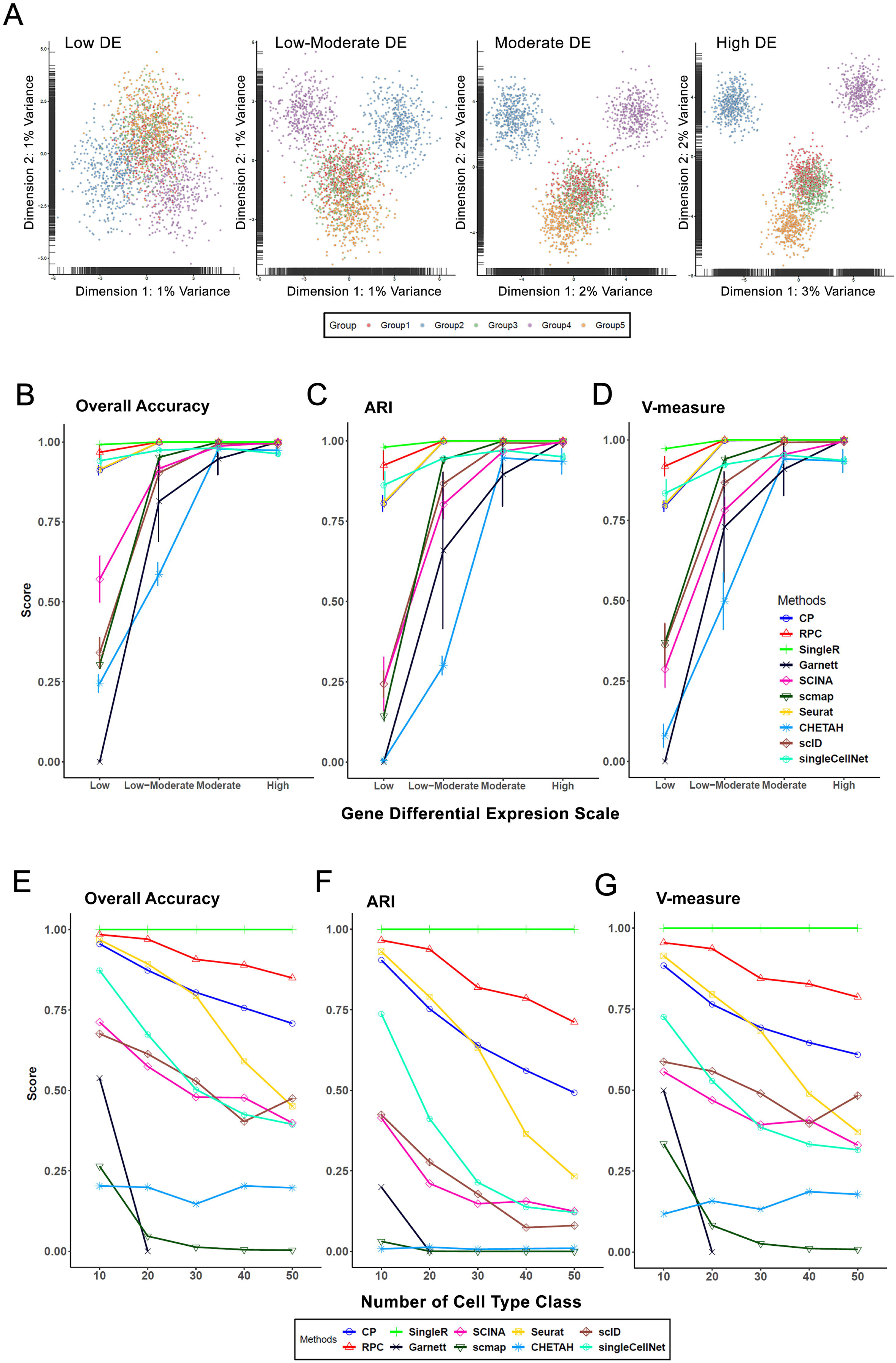
Effect of cell-cell similarity and increased classification labels on annotation tool performance. **(A)** PCA plots of simulation datasets generated by Splatter, each of which is composed of 10,000 genes and 2000 cells, splitting into 5 cell types with equal proportion, and contains the same proportion of differentially expressed genes in each cell type. The datasets differ by changing the magnitude of DE factors for those DE genes to simulate more or less differences between groups. Based on the magnitude of DE factors in five cell groups, we generated 20 datasets with cell groups similarity ranging from low, low-moderate, moderate to high DE (see **Materials and methods**). Colors represent different cell types. True assay (without dropouts) is used as the query and raw assay (with dropout) is used as the reference. **(B-D)** Plots showing three classification metrics to evaluate each annotation method applied to the datasets in (**A**). The x-axis is the DE scale for differential expressed genes in each group, and the y-axis is the metric score. Results are shown as mean+std over 5 repetitions. Line colors and point shapes correspond to different methods. The metrics are: (**B**) overall accuracy, (**C**) adjusted rand index (ARI) and (**D**) v-measure. **(E-G)** Plots illustrating three classification metrics to evaluate each annotation methods applied to five simulation datasets, each of which is composed of an increased number (N) of cell groups (N = 10, 20, 30, 49, 50) with a constant total cell numbers (10,000), gene numbers (20,000), and level of differential expression among cell groups. The x-axis is the number of cell types in each data set, and the y-axis is the metric score. The metrics are: (**E**) overall accuracy, (**F**) adjusted rand index (ARI) and (**G**) v-measure. *Note*: pop_overall: averaged metric scores for all simulations in rare population detection. Low_count: averaged metrics scores for classifying cell types < 1.56% in population. rej_exe_overall: averaged classification metrics score excluding the hold-out group. rej_overall: accuracy of assigning ‘unlabeled’ class to the leave-out group in the query.

### The effect of increased classification labels on annotation performance

The increased cell type classification labels imposed a challenge for some methods in inter and intra-data predictions. We designed five simulation datasets each composed of an increased number (N) of cell groups (N = 10, 20, 30, 40, 50) with a constant total cell numbers, gene numbers, and level of differential expression among cell groups. Similar to the performance that we observed on intra-data and inter-data classification experiments, the increased classification grouping labels lead to dropping accuracy for most methods, except SingleR, which is extremely robust without drop of performance **(Figure 2E-G)**. RPC is consistently ranked 2nd regardless of the cell group numbers. Seurat and CP are ranked the 3nd and 4rd for their robustness before N=30, with small differences in accuracy metrics. However, after N=30, the accuracy of Seurat deteriorates faster and is ranked 4rd instead. The performance issue in Seurat may be due to its susceptibility towards cell-cell similarity. Since we keep a constant differential expression level despite the increased cell grouping labels, more cell types have similar expression profiles and they are more likely to be misclassified. On the other hand, Garnett failed to predict when simulation data set has cell types N>20. Therefore, the simulation study confirms the practical challenge of increased cell labels in multi-label classification for most methods evaluated. SingleR is the most robust method against increased complexity in both real dataset and simulation data evaluations.

### The effect of gene filtering

We also evaluated the stability of annotation methods in inter-dataset classification, by varying the number of query input features. For this purpose, we used the human pancreas data pair (**Table 2**). We randomly down sampled the features from Fluidigm data into 15,000, 10,000 and 5000 input genes, based on the original log count distribution (**Figure 3A**). When the number of features decreases, most methods show decreased metric scores as expected (**Figure 3B**). Seurat and SingleR are the top 2 most robust methods over the decrease of feature numbers, and their ARI scores remain high across all sampling sizes (ARI > 0.9). Again, methods such as Garnett, scID and scmap are more susceptible to low feature numbers, since their performances decrease as the feature number decreases. Therefore, using query data with fewer features than the reference data may affect the prediction performance of those methods. Alternatively, we also downsized the samples by reducing the number of raw reads before alignment and tag counting steps (**Figure 3C**). While most methods show fairly consistent accuracy scores with reduced raw reads as expected, a couple of methods, such as singleCellNet and scID, are perturbed by this procedure (**Figure 3D**).

**Figure 3.**
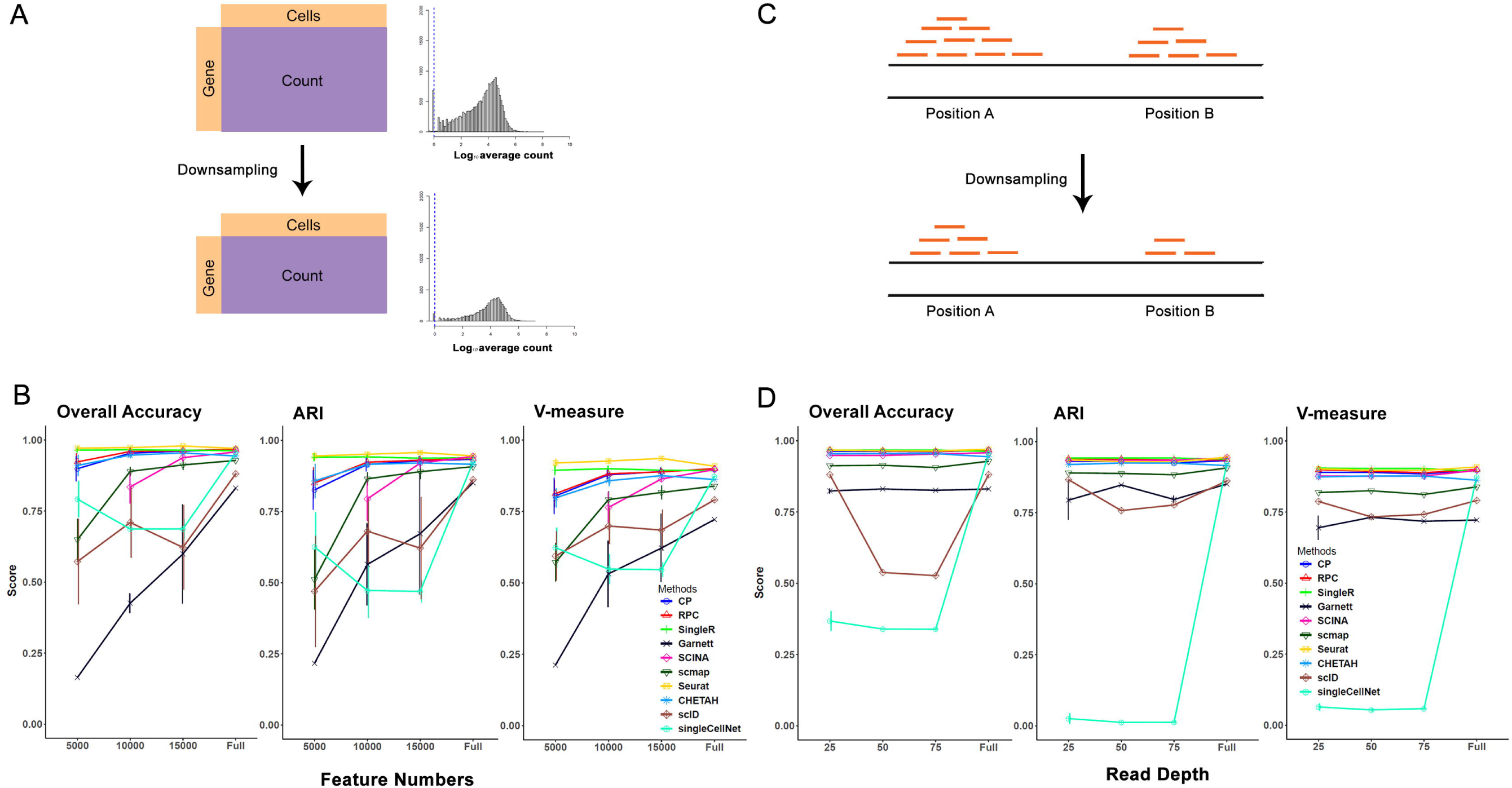
Effects of feature (gene) numbers and read depths on annotation tool performance. **(A)** The features (genes) in the human pancreas-fluidigm dataset are filtered by removing genes that present in less than 3 cells, resulting in 19211 genes. The filtered features (genes) are randomly down sampled into 5000, 10,000 and 15,000 input genes, following the original log count distribution. Such down-sampling was repeated 5 times. **(C)** The BAM file reads in the human pancreas-fluidigm dataset are randomly down sampled into 25%, 50%, 75% of the original read depths. **(B)(D)** Plots depicting the three classification metrics (overall accuracy, adjusted rand index and v-measure) of each method applied to the down sampling approaches in (**A**) and (**C**) respectively. The x-axis is the down sampling size for feature numbers or reads, and the y-axis is the metrics score. Results are shown as mean+std over 5 repetitions. Line colors and point shapes correspond to different methods. SCINA failed when the number of input features reached 5000, thus no point is shown.

### Rare population detection

Identifying rare populations in single cells is a much biologically interesting aspect. We evaluated the inter-dataset classification accuracy per cell population for the top 5 methods based on overall accuracy and adjusted rand index (ARI) (Figure S4): Seurat, SingleR, CP, singleCellNet and RPC (**Figure 1A-B**). We used a mixture of 9 cell populations with a wide variety of percentages (50%, 25%, 12.5%, 6.25%, 3.125%, 1.56%, 0.97%, 0.39%, 0.195%) in ten repeated simulation datasets with different seeds (**Figure 4A**). When the size of the cell population is larger than 50 cells out of 2000 cells, all five methods achieve high cell-type specific accuracy of over 0.8 (**Figure 4B**). However, the classification performances drop drastically for Seurat and singleCellNet when the cell population is 50 or less. On the other hand, most low-performing methods have fluctuated performance and do not perform well in classifying the major cell populations (Figure S4B). Interestingly, bulk-reference based methods such as SingleR, CP and RPC are extremely robust against the size changes of a cell population. They employ averaged profiles as the references and are not susceptible to low cell counts. One challenge for some other single cell methods is that there are not enough cell counts from a low-proportion cell type. Some methods just remove or ignore those cell types in the training phase (such as Garnett), or during alignment (such as Seurat) by their threshold parameters of the algorithms.

**Figure 4.**
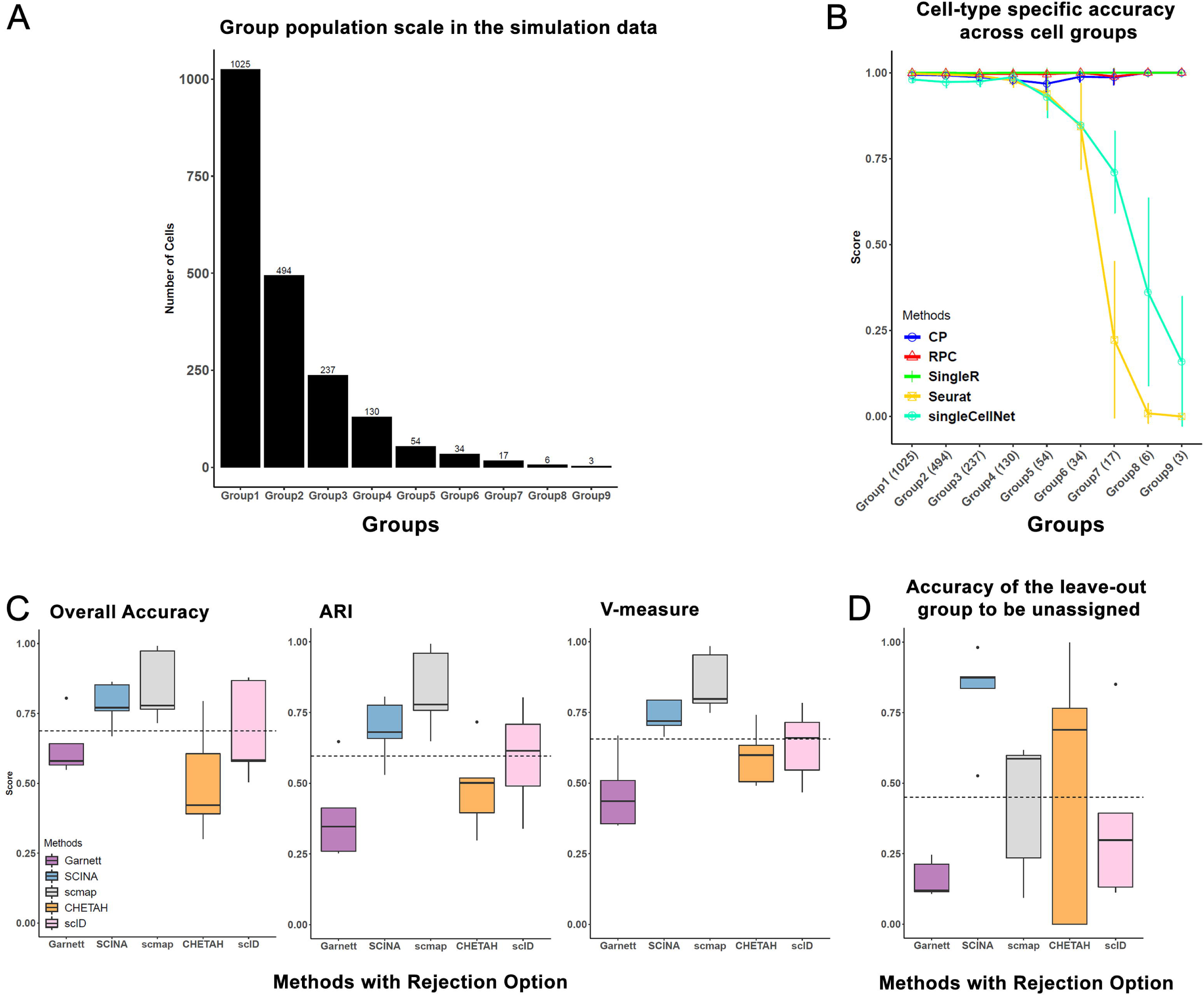
Performance comparison on rare cell type and unknown cell types detection. All datasets are generated by Splatter. **(A)** Cell population distribution of simulation data (10 repeats), composed of 10,000 genes and 2000 cells, split into 9 cell types with proportions of 50%, 25%, 12.5%, 6.25%, 3.125%, 1.56%, 0.97%, 0.39%, 0.195%, respectively. **(B)** Plot illustrating cell-type specific accuracy across 9 cell groups in (**A**), for the five annotation methods that exceed 0.8 in overall accuracy and adjusted rand index (ARI). The x-axis is the cell groups in the descending order for their cell proportions, and the y-axis is the cell-type specific classification score. Results are shown as mean+std over 5 repetitions. **(C)** Performance metrics (overall accuracy, adjusted rand index and v-measure) of another simulation data set, composed of 4000 genes and 2000 cells splitting into 5 cell types. True assay (without dropouts) is used as the reference and the raw assay (with dropout) is used as the query. During each prediction, one cell group is removed from the reference matrix and the query remains intact. The x-axis lists methods with rejection options (e.g. allowing ‘unlabeled’ samples), and the y-axis is the classification metrics score excluding the hold-out group. **(D)** Boxplots showing the accuracy of methods in (**C**), when assigning ‘unlabeled’ class to the leave-out group in the query.

### Unknown population(s) detection

Among the scRNA-seq specific annotation tools, five methods (Garnett, SCINA, scmap, CHETAH, scID) contain the rejection option that allows ‘unassigned’ labels. This is a rather practical option, as the reference data may not contain all cell labels present in the query data. In order to assess how accurate these methods are at labeling ‘unassigned’ cells, we used the scheme of “hold-out one cell type evaluation” on the same simulation dataset pair used in cross-dataset prediction. That is, we remove the signature of one cell type in the reference matrix while keeping the query intact. The evaluation repeated five times for all five cell types. For each method, we measured the average classification accuracies excluding the hold-out group (**Figure 4C**), and the accuracy of assigning unlabeled class to the leave-out group in the query (**Figure 4D**). Among the five methods compared, SCINA, scmap and scID all have metrics scores above the average level of all tools tested for accuracy excluding the hold-out group (**Figure 4C**). However, SCINA has better accuracy in rejecting cell groups existing in the query dataset but not in the reference (**Figure 4D**). Similar results were observed from “hold-out two cell type evaluation” (Figure S5). SCINA has a relatively better balance between overall accuracy in existing cell types and precise rejection in non-existing cell types.

The caveat here, however, is that none of the rejection-enabled methods are among the best performing methods in terms of overall accuracy, stability and robustness to cell type similarities. Since accuracy, stability and robustness are probably more important attributes to assess these methods, the practical guide value based on the results of unknown population detection is limited.

### Time and memory comparison

In order to compare the runtime and memory utilization of the annotation methods, we simulated six data sets each composed of 20,000 genes, with 5 cell types of equal proportion (20%), in total cell numbers of 5000, 10,000, 15,000, 20,000, 25,000, 50,000, respectively (see **Materials and methods**). All methods show increases in computation time and memory usage when the number of cells increases (**Figure 5**). Of the five top-performing methods in the intra-data and inter-data annotation evaluations (**Figure 1**), singleCellNet and CP outperform others on speed (**Figure 5A**). As the dataset size increases beyond 50,000 cells, methods such RPC require a runtime as large as 6 hours. For memory utilization, singleCellNet and CP consistently require less memory than other top performing methods (**Figure 5B**). Notably, the best performing method Seurat (by accuracy) requires memories as large as 100GB, when dataset size increases beyond 50,000 cells, which is significantly larger than most other methods. In all, based on computation speed and memory efficiency, singleCellNet and CP outperform others among the top-class accurate annotation methods.

**Figure 5.**
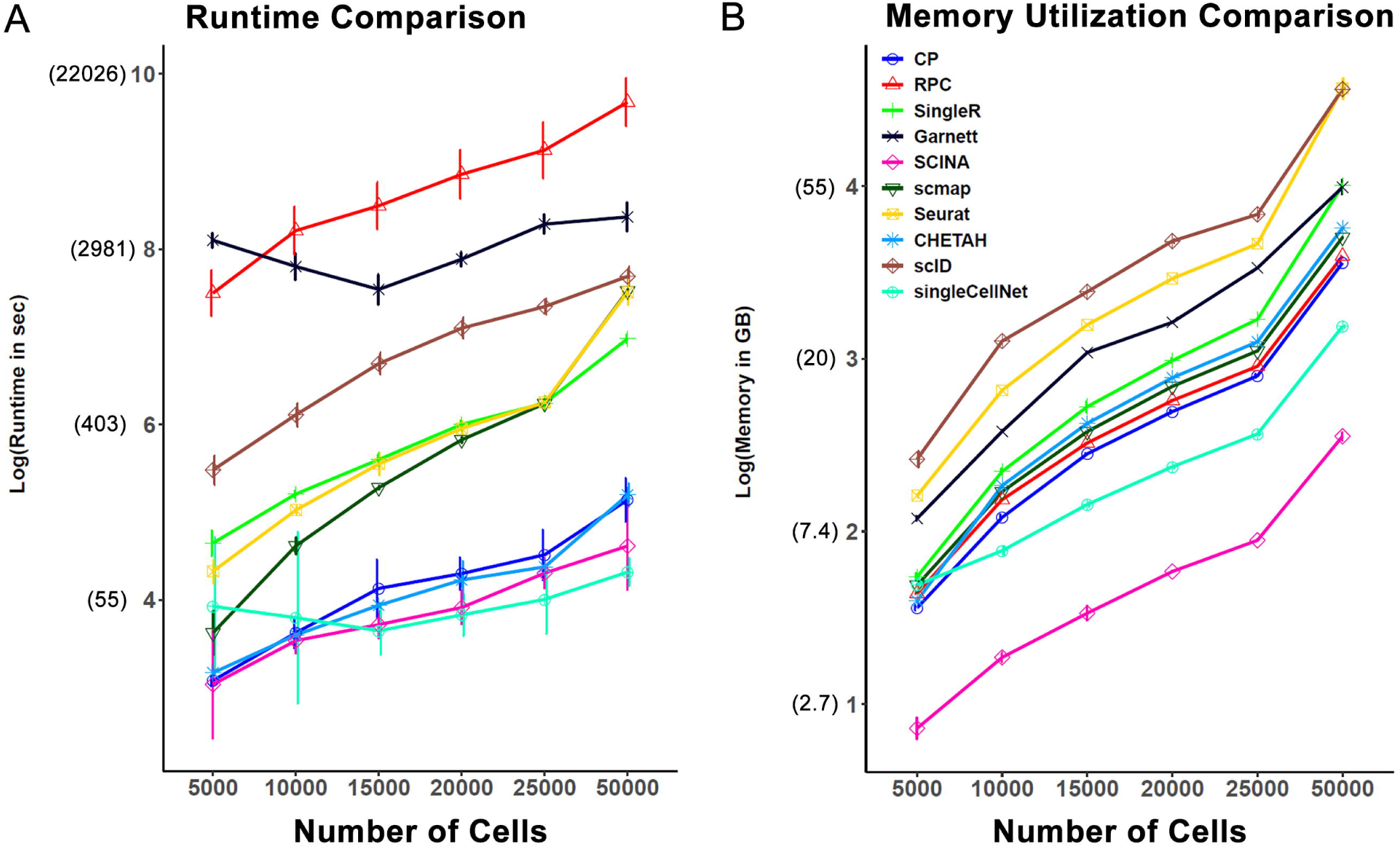
Speed and memory usage comparison. Speed and memory comparison on six pairs of simulation data with increasing numbers of cells (5000, 10,000, 15,000, 20,000, 25,000, 50,000). True assay (without dropouts) is used as the reference and the raw assay (with dropout) is used as the query. Both reference and query contain the same number of cells. Color depicts different annotation methods. **(A)** Natural log of running time (y-axis) vs. cell size (x-axis) over five repetitions in each data point. **(B)** Natural log of peak memory usage (y-axis) vs. cell size (x-axis) over five repetitions in each data point.

## Discussion

In this study, we presented comprehensive evaluations of 10 computational annotation methods in R packages, on single cell RNA-Seq data. Of the 10 methods, 8 of them are designed for single-cell RNA-seq data, and 2 of them are our unique adaptation from methylation-based analysis. We evaluated these methods on 6 publicly available scRNA-seq datasets as well as many additional simulation datasets. We systematically assessed accuracy (through intra-dataset and inter-dataset predictions), the robustness of each method with challenges from gene filtering, cell-types with high similarity, increased cell type classification labels, and the capabilities on rare population detection and unknown population detection, as well as time and memory utilization **(Figure 6)**. In summary, we found that methods such as Seurat, SingleR, CP, RPC and SingleCellNet performed relatively well overall, with Seurat being the best-performing methods in annotating major cell types. Methods such as Seurat, SingleR, RPC and CP are more robust against down-sampling. However, Seurat does have a major drawback at predicting rare cell populations, as well as minor issues at differentiating highly similar cell types and coping with the increased classification labels, while SingleR and RPC are much better in these aspects.

**Figure 6.**
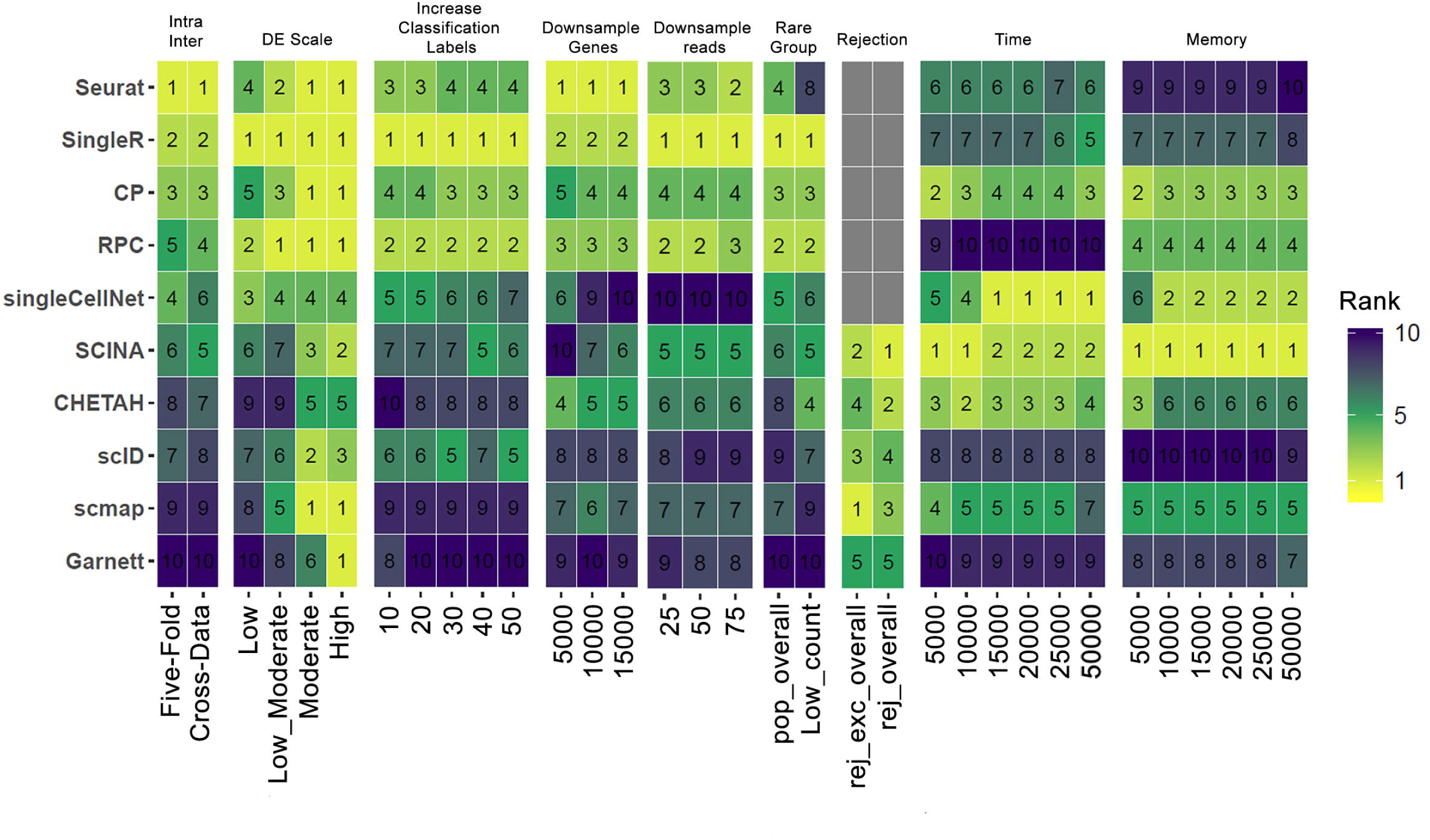
Benchmark Summary. Summary of the classification performance in each evaluation. Each row is a method and each column is evaluations from intra-data and inter-data prediction (Intra-Inter), cell-cell similarity (DE-Scale), increased classification labels, downsampling of genes, downsampling of reads, rare group detection, unknown population detection (rejection), time and memory utilization. The heatmap shows individual method’s rank based on averaged metric scores over overall accuracy, adjusted rand index, and v-measure for each evaluation indicated in the bottom column. Time and memory are ranked by utilization. Grey box indicated that the method does not participate in the evaluation. The methods in the heatmap are arranged in ascending order by their average rank over inter-data and intra-data prediction.

During the preparation of the manuscript, another evaluation paper was published in a special edition of Genome Biology [24]. We, therefore, address the differences between these two studies’ methodologies, before discussing our own findings in detail. First, rather than simply comparing the methods claimed to be “single cell specific”, we uniquely repurpose two methods: Linear Constrained Projection (CP) and Robust Partial Correlations (RPC). Although they were originally developed for DNA methylation data deconvolution, their regression-based principle could be adapted to scRNA-seq supervised/semi-supervised classification. We modified the final regression coefficients as the probability of one specific cell type label, rather than the cell content as in DNA methylation-based deconvolution. As the results indicated, CP and RPC has comparable prediction with SingleR, the overall second best method. This shows the potential of repurposing existing deconvolution methods from another bulk omics analysis. Secondly, for benchmark datasets, we used fewer real experimental datasets. However, we uniquely included many simulated datasets while the other study did not use any. We argue that it is important to have additional simulation datasets, because evaluation based on manually annotated cell-type-specific markers in the experimental data is prone to bias. On the contrary, one can introduce simulation datasets with ‘ground truth’ and unbiasedly assess the tricky issues, such as identifying highly similar cell populations or very rare cell populations. Thirdly, Seurat, the method with the best overall accuracy in our study, is not included in the other study. The high annotation performance of Seurat on intra-data and inter-data predictions in our study, is mostly due to the fact that it’s a classification method using an integrated reference. Its data transfer feature shares the same anchors identification step as the data integration feature. However, unlike data integration, the cell type classification method in Seurat does not correct the query expression data. On top of that, its default setting projects the PCA structure of a reference onto the query, instead of learning a joint structure with CCA [10,21]. This type of methods represents a new trend in single cell supervised classification, evident by a series of scRNA-seq data integration methods (LIGER, Harmony, scAlign *etc* [25–27]). Lastly, we only selected the packages in R with good documentations, as R is still the most popular bioinformatics platform for open-source scRNA-Seq analysis packages (e.g. the arguably most popular method Seurat, which the other study omitted).

Although having slightly lower accuracy metrics scores than Seurat, SingleR and CP still have very excellent performance in intra-data and inter-data prediction, with resilience towards gene filtering and increased complexity in datasets. In addition, SingleR has better performance than Seurat in predicting rare cell populations, dealing with increased cell type classification labels, as well as differentiating highly similar cell types. This advantage of SingleR may benefit from its method and the pseudo-bulk reference matrix. The averaged pseudo-bulk reference profile may potentially remove the variation and noise from the original single cell reference profile, and it can retain the expression profiles of all cell types and is not affected by the low count. SingleR uses pseudo-bulk RNA-seq reference to correlate the expression profiles to each of the single cells in the query data, and uses highly variable genes to find the best fit iteratively. For Seurat, the annotation of the cell labels on query data is informed by the nearest anchor pairs. If two or more cell types have similar profiles, their alignments may overlap which may cause misclassification. Seurat also has some requirements on the minimum number of defined anchor pairs. In the case of rare cell populations, the lack of the neighborhood information makes the prediction difficult. Similar to other study [24], we also found that method that incorporates the prior-knowledge (e.g. Garnett and SCINA) did not improve the classification performance over other methods that do not have such requirements. This prior-knowledge is limited when cell-cell similarity is large. In addition, as the number of cell types increases, the search for the marker genes will become challenging, making these methods even less desirable.

Compared with intra-data prediction, inter-data prediction is more realistic but also more challenging. Technical/platform and batch differences in inter-data prediction may impose major challenges to the classification process, although the tissue and cell type contents are the same. In our study, the CCA batch-correction preprocessing step did not improve the classification accuracy for most methods. Among all experimental data used as the benchmark in this study, PBMC datasets had the worst accuracy results (ARI=0.76 for the best method Seurat). Further inspection of the confusion matrices revealed that the challenges come from distinguishing highly similar cell types, which themselves may have some level of inaccuracy from the original experiments. If the upstream unsupervised clustering methods are not sensitive enough to categorize similar cell populations, this uncertainty may be carried through to the downstream cell annotation steps. This again highlights the potential issue of evaluating the supervised/semi-supervised methods in single cell data, where we are not certain about the ‘ground truth’ of the cell labels to begin with. Recently, some studies used unsupervised classification methods through multi-omics integration, and/or reconstruction of gene regulatory network [28,29], representing a new trend in this area. As the multi-omics technology continues to advance [30], it will be of interest to evaluate these methods, where both multi-omics and pre-defined marker information are available for the same samples.

Overall, we recommend using Seurat for general annotation tasks for cell types that are relatively separable and without rare population identification as the objective. However, for datasets contain cell types with high similarities or rare cell populations, if a reference dataset with clean annotations is available, SingleR, RPC and CP are preferable.

## Materials and methods

### Real data sets

Six real scRNA-seq data sets were downloaded and used for evaluations and validations (**Table 2**). The human pancreatic islet datasets were obtained from the following accession numbers: GEO: GSE85241 (Celseq2) [10,31], GEO: GSE86469 (Fluidigm C1) [10,32]. *The Tabula Muris* datasets Version 2 (10X Genomics and Smart-Seq2) were downloaded from FigShare: https://tabula-muris.ds.czbiohub.org/ [3]. The bead-purified PBMC dataset (10X Genomics) was obtained from the Zheng dataset: https://github.com/10XGenomics/single-cell-3prime-paper, and the PBMC-3K dataset (10X Genomics) was downloaded from https://support.10xgenomics.com/single-cell-gene-expression/datasets [33]. These datasets differ by species, tissue and sequencing protocol. For each of the datasets, we collected both raw counts and cell-type annotations from the corresponding publications, except PBMC-3k, for which the cell-type annotations were obtained through the standard single cell RNA-seq analysis and classified using cell-type-specific marker genes. The extracted cell-type annotations for each dataset were used as the ground truth for evaluations (Table S1).

#### Data cleaning

Datasets were paired in groups by tissue type (**Table 2**). Within a pair, we used the data generated by Fluorescence-activated cell sorting (FACS) sorted method as reference data. Both reference data and query data were further processed to make sure the cell types in reference data are larger or equal to the cell types in the query data. When necessary, the query data were down sampled following the original cell type count distribution. For the two Tabula Muris (TM-Full) datasets from 10X and Smart-Seq2 platforms, which contain a large number of cell types (32 and 37 cell types, respectively), we took a subset of cells from lung tissue and created two TM-Lung datasets that have fewer cell types, 8 for 10X and 10 for Smart-Seq2 platform, respectively. As a result, we have four pairs of experimental datasets: PBMC cell pair with PBMC-sorted-ref and PBMC-3K- query; pancreas cell pair with pancreas-celseq2-ref and pancreas-fluidigm-query; TM-Full pair with TM-Full-smartseq2-ref and TM-Full-10X-query; TM-Lung pair with TM-Lung-smartseq2- ref and TM-Lung-10x-query.

#### Data downsampling

To explore the effects of different feature numbers and read depths on the performance of tools, we randomly down sampled features (genes) from human pancreas-Fluidigm dataset into 5000, 10,000 and 15,000 input genes, following the original log count distribution. We repeated five times for each downsampling scheme. Alternatively, we also down sampled the reads into 25%, 50%, 75% of the original read depths (with 2 repetitions) using *samtools* on BAM files, and then realigned following the method provided by the original manuscript [32].

### Simulated Data Sets

We simulated a dataset using Splatter, with 4000 genes and 2000 cells (Splatter parameters, dropout.shape=-0.5, dropout.mid=1), and then split each dataset into 5 cell groups with proportions 10%, 30%, 30%, 10% and 20%. In addition, we also generated three additional simulation sets to evaluate the robustness of tools. In the first set, we generated 10 simulation datasets each has 10,000 genes and 2,000 cells (use Splatter parameters dropout.shape=-0.5, dropout.mid=1, 10 different seeds), and then split each into 9 cell groups with proportions 50%, 25%, 12.5%, 6.25%, 3.125%, 1.56%, 0.97%, 0.39%, 0.195%, respectively. The second set contains 20 simulation datasets, each composed of 10,000 genes and 2,000 cells splitting into 5 cell types with equal proportions. These datasets have the same set of differentially expressed (DE) genes, but differ by the magnitude of DE factors (*de*.*facScale* parameter in Splatter). We simulated each DE scale five times with five different seeds. The DE scales and the parameterizations are: low: de.facScale = c(0.1, 0.3, 0.1, 0.3, 0.2); low-moderate: de.facScale = c(0.3, 0.5, 0.3, 0.5, 0.4); moderate: de.facScale = c(0.5, 0.7, 0.5, 0.7, 0.6); high: de.facScale = c(0.7, 0.9, 0.7, 0.9, 0.8). The third set contains five simulation datasets each composed of an increased number (N) of cell groups (N = 10, 20, 30, 49, 50) with a constant total cell numbers (10,000), gene numbers (20,000), and level of differential expression among cell groups. Each simulation dataset contains two paired assays. The true assay without dropouts was used as the reference and the raw assay with dropout mask was used as the query.

### Data Preprocessing

#### Cell and gene filtering

We filtered out cells for which fewer than 200 genes were detected and any genes that were expressed in fewer than 3 cells.

#### Normalization

For the annotation tools that require a normalized count matrix as input, we performed log-normalization using a size factor of 10,000.

#### Pseudo-bulk reference matrix

For the annotation tools that use bulk rather than single-cell expression profiles as reference, we took the average of the normalized count of each cell type group and made a pseudo-bulk RNA-seq reference.

#### Marker genes selection

Some classification tools (SCINA and Garnett) require cell-type specific marker as the input. When such marker information is neither provided by the corresponding tools nor retrievable by public research, we extract them from the reference data by performing differential expression analysis using Wilcoxon rank sum test (*FindAllMarkers* function from Seurat with parameters only.pos = TRUE, min.pct = 0.25 and logfc.threshold = 0.25). Wilcoxon rank sum test is the most common nonparametric test for a difference in mean expression between cell groups. The top 10 ranked marker genes for each cell type were used as the input for the corresponding tools.

### Supervised/Semi-supervised Annotation Methods

We only considered pre-printed or published methods with detailed documentation on installation and execution. We excluded any methods that required extensive running time, and where we were unable to customize the reference dataset, or random and inconsistent predictions were produced. In the end, ten cell annotation methods, publicly available as R packages, were evaluated in this study. This includes eight methods (Seurat, scmap, SingleR, CHETAH, SingleCellNet, scID, Garnett, and SCINA) commonly used to annotate scRNA-seq data. In addition, to investigate the potential to repurpose deconvolution methods for other bulk omics analysis, we also included and modified two methods originally designed for bulk DNA methylation that use a different type of algorithms not yet reported in scRNA-seq specific tools: Linear Constrained Projection (CP) and Robust Partial Correlations (RPC).

All parameters were set to default values following the author’s recommendations or the respective manuals (**Table 1**). For methods that allow “unknown” assignments (scmap, CHETAH, scID, Garnett, and SCINA), we modified the parameter to force assignments where possible (except for the evaluations where unknown assignments were allowed).

#### Adaptation of CP and RPC methods for scRNA-Seq analysis

In order to accommodate the methylation-based methods for scRNA-seq data, we made some modifications. In original papers, both RPC and CP model the methylation profile of any given sample as a linear combination of a given set of reference profiles representing underlying cell-types present in the sample. Assume the number of underlying cell-types to be **C**, and each cell type has a profile **b**_**c**_ that constitutes the signature matrix **H** [34–36]. **Let y** be the profile of a given sample and w_c_ be the weight estimation of cellular proportion of each cell type, and the underlying model becomes:

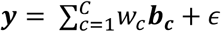

Both methods assume that reference profiles contain the major cell-types present in the sample **y** and sum of weights equal to 1. RPC estimates the weight coefficients using robust multivariate linear regression or robust partial correlation, while CP uses a quadratic programming technique known as linear constrained protection to estimate the weights [37].

In the modified version, we first converted the single cell RNA-seq reference data into pseudo-bulk RNA-seq data matrix by taking the average of the normalized count of each cell type group. Then we took the subset of pseudo-bulk RNA-seq data by keeping *n* features that exhibited high cell-to-cell variations across **C** distinct cell types in the reference dataset, and had a small condition number below 3 as the signature matrix **H** [34]. **We set the highly variable genes to 2000, using *FindVaraibleFeatures* function from Seurat (Figure S6). We let y** be the profile of a given single cell from the query data with the same 2000 genes from the signature matrix **H**. While applying both algorithms, we treated the estimated weight for each cell type as the probability and the cell type with the highest weight was the identity of the corresponding single cell sample in the query data. This conversion is based on the fact that **y** no longer represents averages over many different cell types, but only expression profile from only one cell type (since we have single cell data).

### Benchmarking

#### Five-fold cross validation and cross-dataset prediction

For each dataset in four pairs of the real experimental datasets mentioned above, we used a 5-fold cross validation where the four-fold data were used as the reference and the remaining one-fold as the query. For the cross-dataset prediction, in addition to the four pairs of real datasets, we used simulation datasets containing true assay (without dropouts) as the reference and raw assay (with dropout mask) as the query.

In order to evaluate whether batch correction and data integration benefit the classification performance, for each pair of real dataset, we aligned both reference and query dataset using CCA [10,21] from the Seurat data integration function. Then we separated the aligned datasets and performed the cross-dataset evaluation again.

#### Performance evaluation on the effect of feature numbers and read depths

To investigate the robustness of different methods with regards to feature numbers and read depths, we used the down-sampled human pancreas Fluidigm data set as described in the data downsampling section. In such evaluation, the human pancreas Celseq2 dataset was used as the reference and the down-sampled human pancreas Fluidigm dataset was used as a query.

#### Performance evaluation with effect of differential expression (DE) scale among cell groups

In this assessment, we used 20 simulation data sets containing the same DE gene set but differing only by DE factors as described earlier in the Simulated Data Sets section. Each simulation data set contains two paired assays. The true assay (without dropouts) was used as the reference and the raw assay (with dropout mask) was used as the query.

#### Performance evaluation on the effect of increased classification labels

In this evaluation, we designed five simulation data sets, each composed of an increased number (N) of cell groups (N=10, 20, 30, 40, 50) with a constant total cell numbers, gene numbers, and level of differential expression among cell groups. Each simulation data set contains two paired assays. The true assay (without dropouts) was used as the reference and the raw assay (with dropout mask) was used as the query.

#### Rare and unknown population detection

Each of the 10 simulation data sets in the rare population detection evaluation was composed of 10, 000 genes and 2000 cells splitting into 9 cell types with proportions 50%, 25%, 12.5%, 6.25%, 3.125%, 1.56%, 0.97%, 0.39%, 0.195%. The simulation dataset in the unknown population detection evaluation was composed of 4000 genes and 2000 cells splitting into 5 cell types. We used the scheme of “hold-out one cell type evaluation” to evaluate prediction on the unknown population, that is, removing the signature of one cell type in the reference matrix while predicting the query. During each prediction, one cell group was removed from the reference matrix and the query remained intact. We repeated the evaluation five times for all five cell types. We additionally employed a “hold-out two cell type” experiment, in which we removed signatures of any combination of two cell types in the reference matrix while keeping the query intact. The evaluation was repeated ten times for all ten different combinations. Similarly, for each simulation data set, the true assay (without dropouts) was used as the reference and the raw assay (with dropout mask) was used as the query.

#### Runtime and Memory Assessment

In order to compare the computational runtime and memory utilization of annotation methods, we simulate six datasets, with total cell numbers of 5000, 10,000, 15,000, 20,000, 25,000, and 50,000, respectively, each composed of 20,000 genes, splitting into 5 cell types with the equal proportion. The true assay (without dropouts) was used as the reference and the raw assay (with dropout mask) was used as the query. Each execution was performed in a separate R session in our lab server (4 nodes (Dell PowerEdge C6420) of 2 X Intel(R) Xeon(R) Gold 6154 CPU @ 3.00GHz, 192GB RAM, one node (Dell Poweredge R740) with 2 X Xeon(R) Gold 6148 CPU @ 2.40GHz, 192 GB RAM, and two 16GB Nvidia V100 GPUs) with Slurm job scheduler. One processor and 100GB memory were reserved for each job. From the job summary, we collected ‘Job Wall-clock time’ and ‘Memory Utilized’ for evaluation. We ran each method on each dataset five times to estimate the average computation time.

### Evaluation Criteria

The prediction results of the methods are evaluated using three different metrics: overall accuracy, adjusted rand index, and V-measure. We used three different metrics to avoid possible bias in evaluating the performance. The detailed explanations on these metrics were described earlier [22,38,39]. Briefly, *Overall accuracy* is the percent agreement between the predicted label and the true label. *Adjusted rand index* (ARI) is the ratio of all cell pairs that are either correctly classified together or correctly not classified together, among all possible pairs, with adjustment for chance. *V-measure* is computed as the harmonic mean of distinct homogeneity and completeness score. In specific, homogeneity is used to assess whether each predicted cell type groups contains only members of a single class, while completeness is used to assess whether all members of a given class are assigned to the same predicted cell label.

## Code Availability

All the codes and data are available at: https://github.com/qianhuiSenn/scRNA_cell_deconv_benchmark.

## Authors’ Contributions

LG, and QH envisioned this project. QH implemented the project and conducted the analysis with help from YL and YD. QH and LG wrote the manuscript. All authors have read and agreed on the manuscript.

## Acknowledgements

This research was supported by grants K01ES025434 awarded by NIEHS through funds provided by the trans-NIH Big Data to Knowledge (BD2K) initiative (www.bd2k.nih.gov), R01 LM012373 and R01 LM012907 awarded by NLM, R01 HD084633 awarded by NICHD to L.X. Garmire.

## Supplementary material

### Supplementary Figures

**Supplementary Figure 1.**
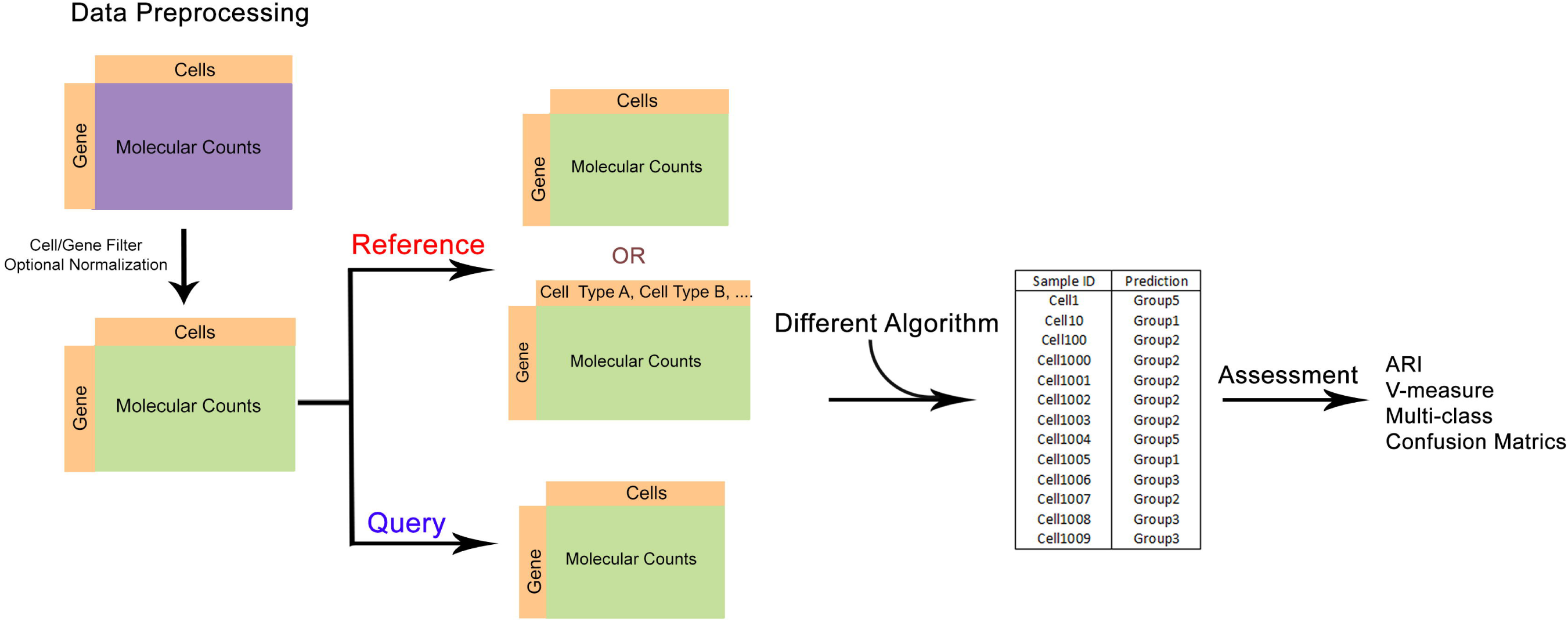
Benchmark Workflow. Illustration of the workflow for this study consists of 1) preprocessing 2) prediction 3) evaluations.

**Supplementary Figure 2.**
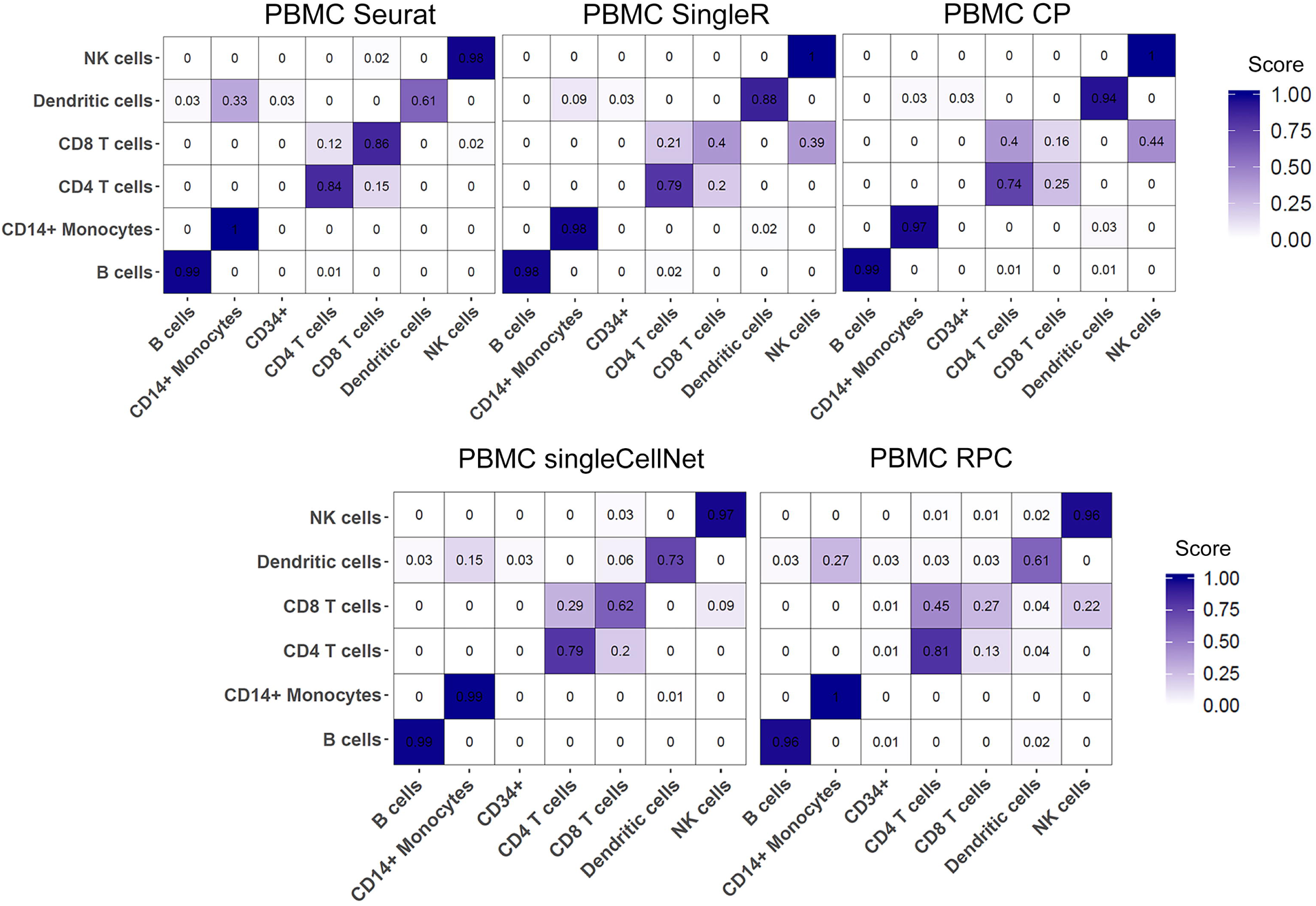
Cell-type specific accuracy for top 5 performing methods on PBMC cross-dataset prediction. Confusion matrix of cell-type specific accuracy for PBMC inter-dataset predictions among top performing annotation methods (Seurat, SingleR, CP, singleCellNet, RPC). The x-axis is the predicted label from each algorithm, and the y-axis is the true label in the query data.

**Supplementary Figure 3.**
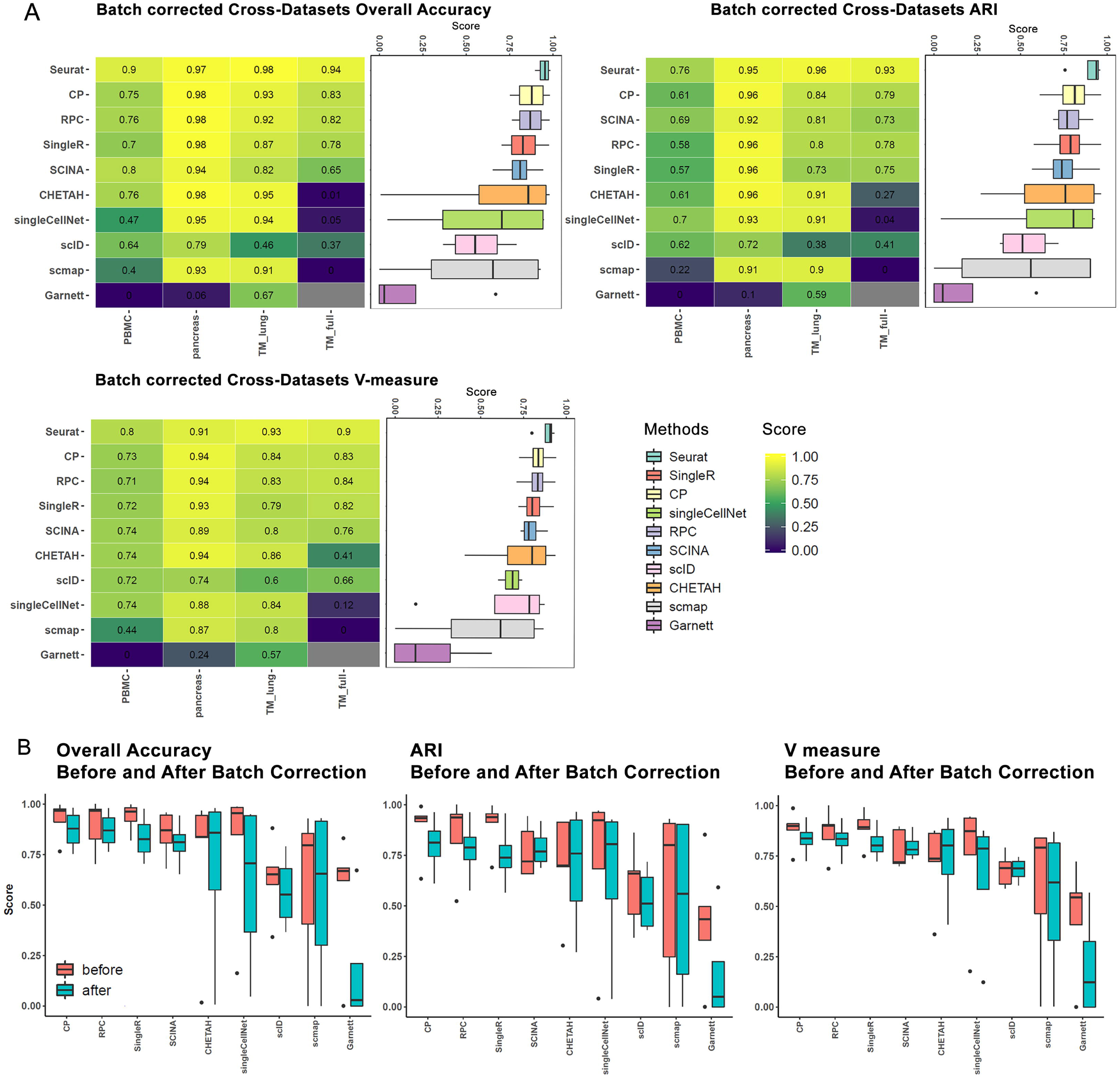
Inter-data prediction using aligned reference and query matrix. For each of the four pairs of experimental data used in cross-data evaluation, we aligned both reference and query dataset using CCA from the Seurat data integration function. Then we separated the aligned datasets and performed the cross-dataset evaluation again. **(A)** Inter-data accuracy comparison, shown as heatmaps of three classification metrics (overall accuracy, adjusted rand index (ARI), and v-measure). **(B)** Boxplots illustrating the averaged metrics scores before and after alignment for each method. The x-axis is the methods, and the y-axis is the classification metrics score.

**Supplementary Figure 4.**
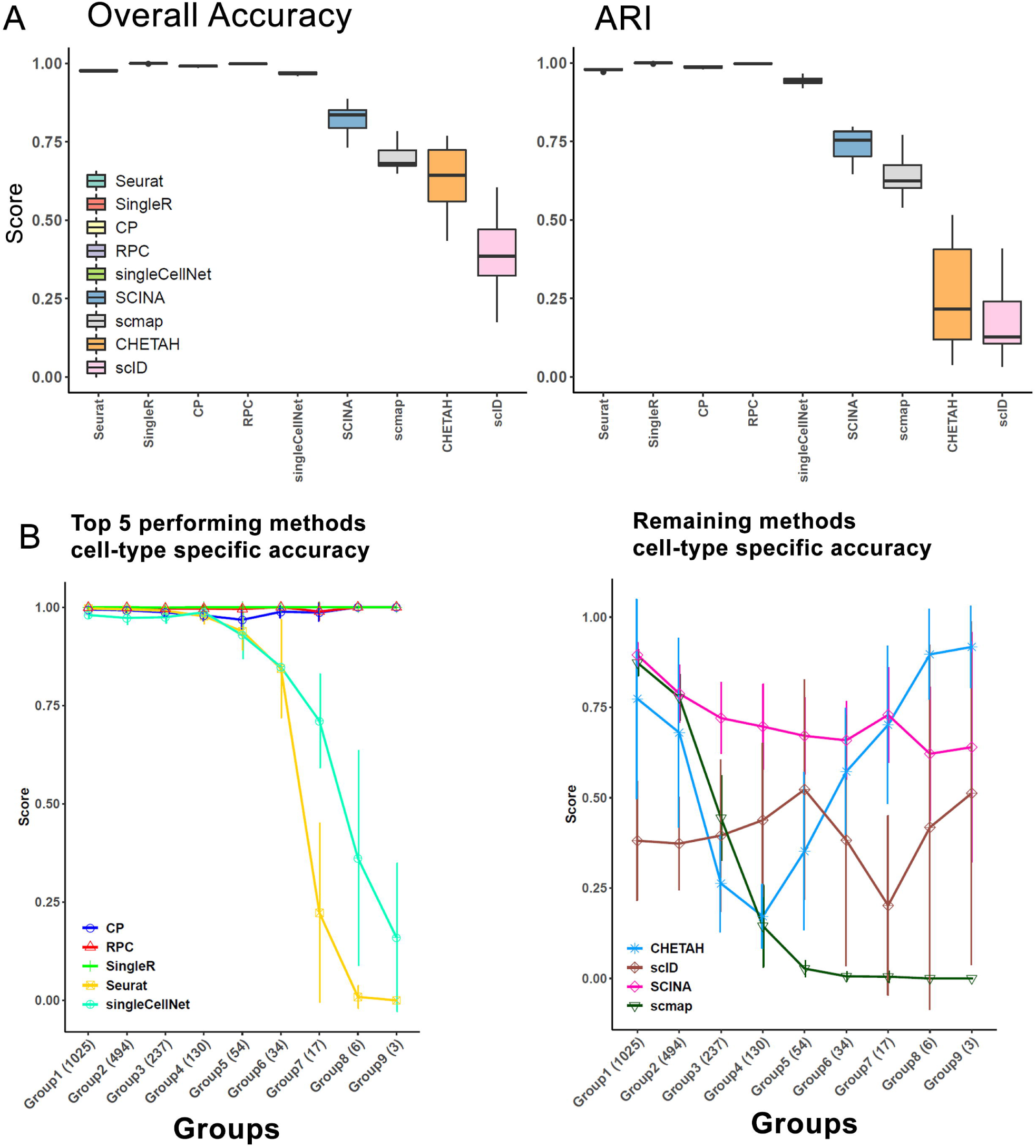
Rare population detection evaluation for remaining 5 methods. **(A)** Boxplots illustrating the averaged overall accuracy and adjusted rand index over all the rare population detection simulation data. The x-axis is the methods evaluated, and the y-axis is the metric score. **(B)** Rare population detection results for the five methods with lower overall accuracy and ARI. The x-axis is the cell groups in the descending order for their cell proportions, and the y-axis is the cell-type specific classification score.

**Supplementary Figure 5.**
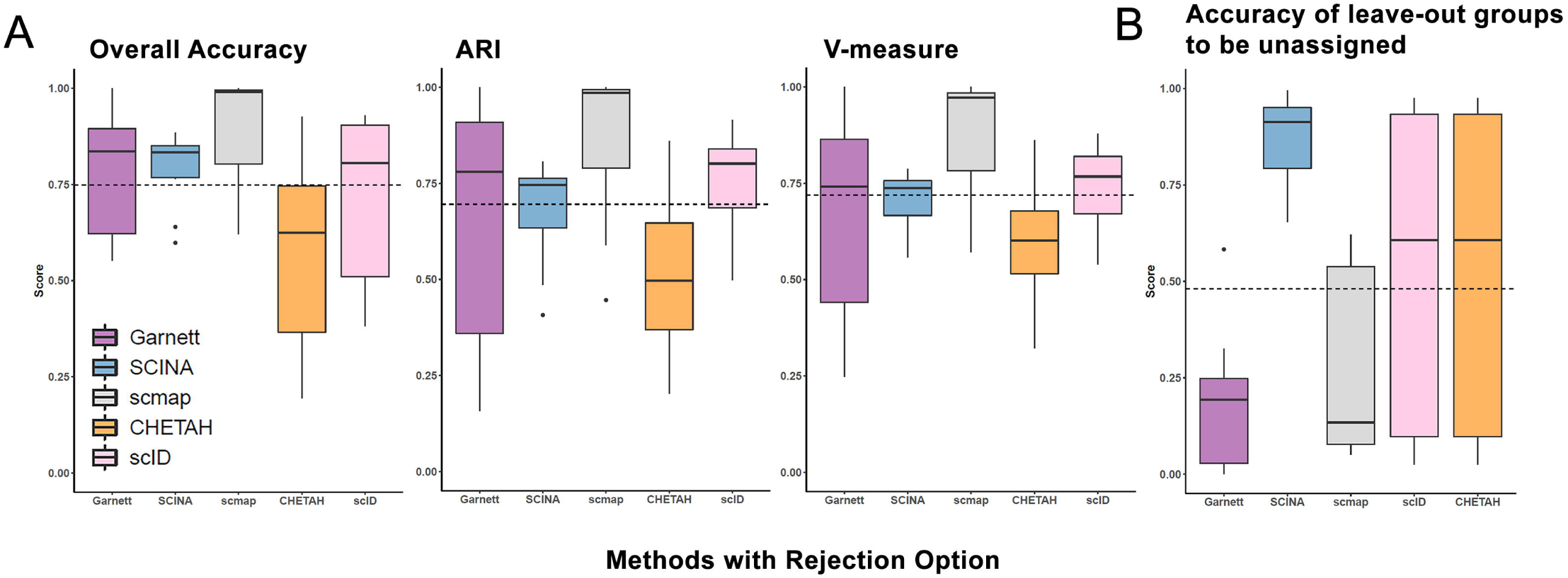
Hold-out two cell type rejection evaluation. “Hold-out two cell type” experiment was performed on the same simulation dataset pair used in cross-dataset prediction. In this experiment, signatures of any combination of two cell types were removed in the reference matrix while keeping the query intact. The evaluation was repeated ten times for all ten different combinations. **(A)** The x-axis lists methods with rejection options (e.g. allowing ‘unlabeled’ samples), and the y-axis is the classification metrics score excluding the hold-out groups. **(B)** Boxplots showing the accuracy of methods in (**A**), when assigning ‘unlabeled’ class to leave-out groups in the query.

**Supplementary Figure 6.**
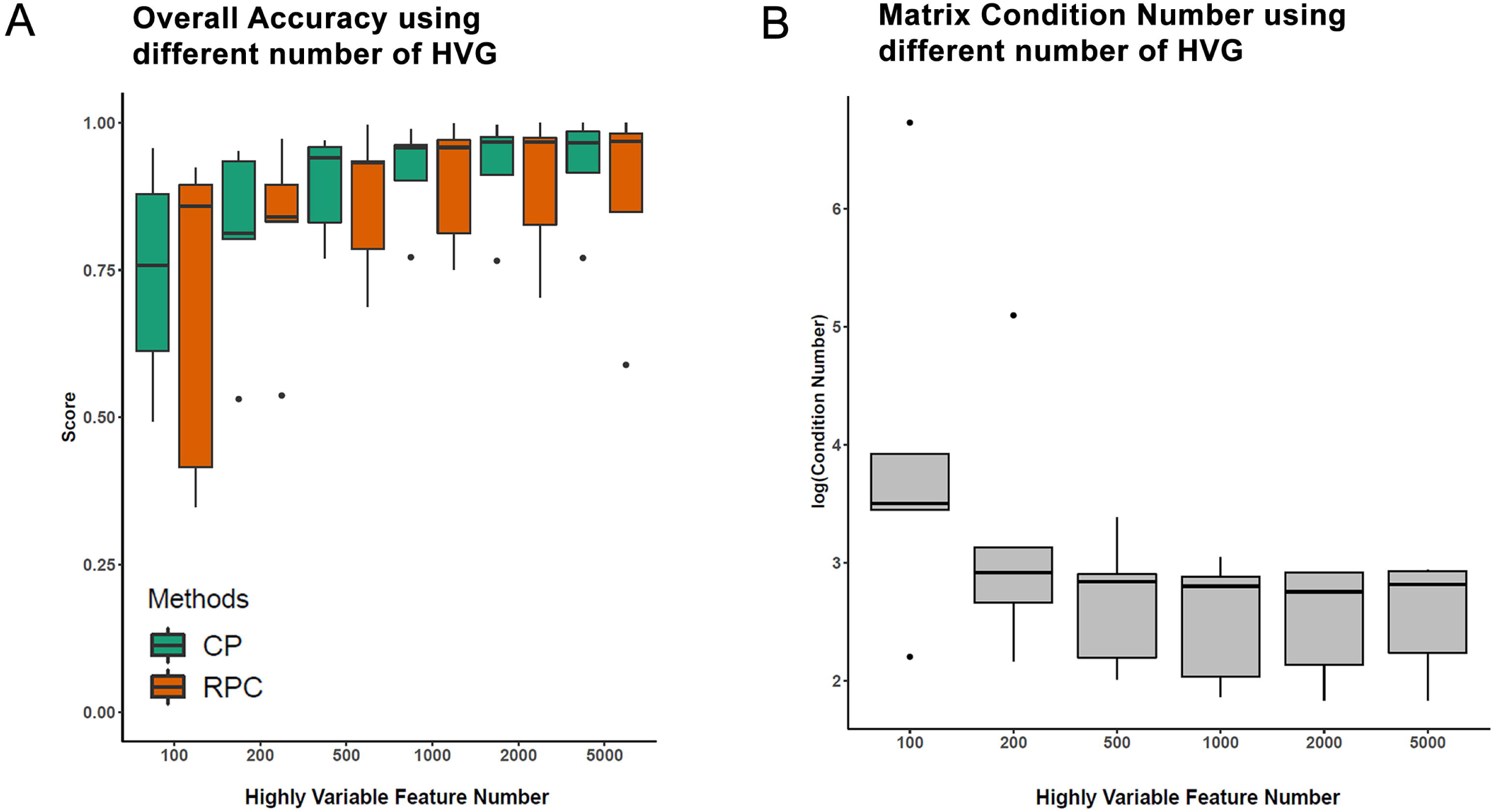
The optimal number of highly variable genes (HVG) to be used in CP and RPC algorithms. The highly variable genes are identified from reference dataset and ranked by standardized variance from mean-variance feature selection methods with variance-stabilizing transformation. **(A)** The boxplot depicts the overall accuracy averaged over five pairs of inter-dataset predictions (pbmc, pancreas, tabula-Full, tabula-Lung, and simulation) with the top 100, 200, 500, 1000, 2000, and 5000 highly variable genes as input features for CP and RPC methods. The x-axis is the number of highly variable features, and the y-axis is the overall accuracy. Methods are reflected by different box colors. **(B)** The boxplot represents the condition number of the pseudo-bulk reference matrix averaged over four combinations of cross-dataset predictions with the top 100, 200, 500, 1000, 2000, and 5000 highly variable genes as input features. The x-axis is the number of highly variable features, and the y-axis is the condition number.

### Supplementary Tables

**Supplementary Table 1.** Composition of cell-types in each real dataset.

| **PBMC-Sorted** | **PBMC-3K** | **Pancreas-celseq2** | **Pancreas-Fluidigm** | **Tabula Muris-Lung-smartseq2** | **Tabula Muris-Lung-10X** | **Tabula Muris-Full-smartseq2** | **Tabula Muris-Full-10X** |
| --- | --- | --- | --- | --- | --- | --- | --- |
| B cells (10,084) | B cells (342) | Acinar<br>(274) | Acinar<br>(21) | B cells<br>(57) | B cells<br>(140) | B cells<br>(2029) | B cells<br>(5615) |
| CD14+<br>Monocytes<br>(2,465) | CD14+<br>Monocytes<br>(2,465) | Activated<br>stellata<br>(90) | Activated<br>stellata<br>(16) | Ciliated<br>columnar cell<br>(25) | - | Basal cell<br>(1340) | Basal cell<br>(27) |
| CD34+<br>(6,312) | - | Alpha<br>(843) | Alpha<br>(239) | Classical<br>monocyte<br>(90) | Classical<br>monocyte<br>(4) | Basal cell of epidermis<br>(1648) | Basal cell of<br>epidermis<br>(2020) |
| CD4+ T cells<br>(42,166) | CD4+ T cells<br>(479) | Beta<br>(445) | Beta<br>(258) | Leukocyte<br>(35) | Leukocyte<br>(9) | Basophil<br>(25) | - |
| CD8+ T cells<br>(22,138) | CD8+ T cells<br>(308) | Delta<br>(203) | Delta<br>(25) | Lung endothelia<br>cell<br>(693) | Lung<br>endothelia cell<br>(24) | Bladder cell<br>(695) | Bladder cell<br>(192) |
| Dendritic cells<br>(99) | Dendritic<br>cells<br>(33) | Ductal<br>(258) | Ductal<br>(36) | Monocyte<br>(65) | - | Bladder urothelial cell<br>(683) | Bladder urothelial<br>cell<br>(141) |
| NK cells<br>(8,385) | NK cells<br>(155) | Endothelial<br>(21) | Endothelial<br>(14) | Myeloid cell<br>(85) | Myeloid cell<br>(2) | Blood cell<br>(206) | Blood cell<br>(153) |
| - | - | Epsilon<br>(4) | Epsilon<br>(1) | Natural killer<br>cell<br>(37) | Natural killer<br>cell<br>(113) | Cardiac muscle cell<br>(133) | - |
| - | - | Gamma<br>(110) | Gamma<br>(18) | Stromal cell<br>(423) | Stromal cell<br>(888) | Ciliated columnar cell<br>(25) | - |
| - | - | Macrophage<br>(15) | Macrophag<br>e<br>(1) | T cell<br>(53) | T cell<br>(123) | Classical monocyte<br>(90) | Classical monocyte<br>(4) |
| - | - | Mast<br>(6) | Mast<br>(3) | - | - | DN1 thymic pro-T cell<br>(32) | - |
| - | - | Quiescent<br>stella | Quiescent<br>stella | - | - | Endocardial cell<br>(165) | Endocardial cell<br>(1) |
|  |  | (12) | (1) |  |  |  |  |
| - | - | Schwann | Schwann | - | - | Endothelial cell | Endothelial cell |
|  |  | (4) | (5) |  |  | (3319) | (971) |
| - | - | - | - | - | - | Endothelial cell of<br>hepatic sinusoid | - |
|  |  |  |  |  |  | (182) |  |
| - | - | - | - | - | - | Epithelial cell | Epithelial cell |
|  |  |  |  |  |  | (201) | (99) |
| - | - | - | - | - | - | Fibroblast | Fibroblast |
|  |  |  |  |  |  | (2189) | (4) |
| - | - | - | - | - | - | Granulocyte | Granulocyte |
|  |  |  |  |  |  | (761) | (73) |
| - | - | - | - | - | - | Granulocytopoietic cell | Granulocytopoietic<br>cell |
|  |  |  |  |  |  | (221) | (14) |
| - | - | - | - | - | - | Hematopoietic precursor<br>cell | Hematopoietic<br>precursor cell |
|  |  |  |  |  |  | (265) | (24) |
| - | - | - | - | - | - | Hepatocyte<br>(391) | Hepatocyte<br>(374) |
| - | - | - | - | - | - | Immature B cell<br>(344) | Immature B cell<br>(3) |
| - | - | - | - | - | - | Immature T cell<br>(1337) | Immature T cell<br>(222) |
| - | - | - | - | - | - | Keratinocyte<br>(330) | Keratinocyte<br>(1035) |
| - | - | - | - | - | - | Kidney collecting duct<br>epithelial cell<br>(121) | Kidney collecting<br>duct epithelial cell<br>(24) |
| - | - | - | - | - | - | Late pro-B cell<br>(306) | Late pro-B cell<br>(14) |
| - | - | - | - | - | - | Leukocyte<br>(683) | Leukocyte<br>(19) |
| - | - | - | - | - | - | Luminal epithelial cell<br>of mammary gland | Luminal epithelial<br>cell of mammary<br>gland |
|  |  |  |  |  |  | (578) | (35) |
| - | - | - | - | - | - | Lung endothelial cell | Lung endothelial cell |
|  |  |  |  |  |  | (693) | (24) |
| - | - | - | - | - | - | Macrophage | Macrophage |
|  |  |  |  |  |  | (395) | (208) |
| - | - | - | - | - | - | Mesenchymal cell | Mesenchymal cell |
|  |  |  |  |  |  | (830) | (5200) |
| - | - | - | - | - | - | Mesenchymal stem cell | Mesenchymal stem cell |
|  |  |  |  |  |  | (499) | (169) |
| - | - | - | - | - | - | Monocyte | Monocyte |
|  |  |  |  |  |  | (331) | (42) |
| - | - | - | - | - | - | Myeloid cell | Myeloid cell |
|  |  |  |  |  |  | (1208) | (2) |
| - | - | - | - | - | - | Natural killer cell | Natural killer cell |
|  |  |  |  |  |  | (171) | (142) |
| - | - | - | - | - | - | Skeletal muscle satellite cell | Skeletal muscle satellite cell |
|  |  |  |  |  |  | (540) | (11) |
| - | - | - | - | - | - | Stromal cell | Stromal cell |
|  |  |  |  |  |  | (863) | (1153) |
| - | - | - | - | - | - | T cell | T cell |
|  |  |  |  |  |  | (793) | (1985) |

*Note:* Values are indicated as *Cell Type (Cell count)*

